# Synthesis of higher order feature codes through stimulus-specific supra-linear summation

**DOI:** 10.1101/2020.06.24.169359

**Authors:** Evan H. Lyall, Daniel P. Mossing, Scott R. Pluta, Amir Dudai, Hillel Adesnik

**Affiliations:** Biophysics graduate program; Department of Molecular and Cell Biology; Department of Biology, Purdue University; The Edmond and Lily Safra Center for Brain Sciences and The Life Sciences Institute, The Hebrew University of Jerusalem, Israel; The Helen Wills Neuroscience Institute, University of California, Berkeley

## Abstract

How cortical circuits build representations of complex objects is poorly understood. The massive dimensional expansion from the thalamus to the primary sensory cortex may enable sparse, comprehensive representations of higher order features to facilitate object identification. To generate such a code, cortical neurons must integrate broadly over space, yet simultaneously obtain sharp tuning to specific stimulus features. The logic of cortical integration that may synthesize such a sparse, high dimensional code for complex features is not known. To address this question, we probed the integration and population coding of higher order stimuli in the somatosensory and visual cortices of awake mice using two-photon calcium imaging across cortical layers. We found that somatosensory and visual cortical neurons sum highly specific combinations of sensory inputs supra-linearly, but integrate other inputs sub-linearly, leading to selective responses to higher order features. This integrative process generates a sparse, but comprehensive code for complex stimuli from the earliest stages of cortical processing. These results from multiple sensory modalities imply that input-specific supra-linear summation may represent a widespread cortical mechanism for the synthesis of higher order feature codes. This new mechanism may explain how the brain exploits the thalamocortical expansion of dimensionality to encode arbitrary complex features of sensory stimuli.

## Introduction

Cortical neurons must encode diverse features of sensory stimuli to give rise to unique sensory percepts. By integrating inputs within their receptive fields, cortical neurons extract local features of the sensory scene(Bensmaia et al., 2008; Hubel and Wiesel, 1959; Priebe and Ferster, 2012). By integrating inputs beyond their receptive fields, cortical neurons can extract more global sensory features, such as contour and shape(Kapadia et al., 1995; Kourtzi et al., 2003; Li and Gilbert, 2002; Stettler et al., 2002), which may be critical for object identification. Local summation is exemplified by primary visual cortical neurons, which extract stimulus orientation by linearly summing specific combinations of spatially offset thalamocortical inputs within their receptive fields(Hubel and Wiesel, 1962). Global (or ‘contextual’) summation is far less understood; except in rare instances(Ego-Stengel et al., 2005; Estebanez et al., 2016; Hirata and Castro-Alamancos, 2008; Ramirez et al., 2014; Shimegi et al., 1999), adding stimuli outside neurons’ receptive fields yields sub-linear summation, or even suppression(Adesnik et al., 2012; Angelucci et al., 2017; Arabzadeh et al., 2003; Bair et al., 2003; Blakemore and Tobin, 1972; Brumberg et al., 1996; Cavanaugh et al., 2002; Dipoppa et al., 2018; Ego-Stengel et al., 2005; Fanselow and Nicolelis, 1999; Gilbert, 1977; Kato et al., 2017; Mirabella et al., 2001a; Ozeki et al., 2009; Shushruth et al., 2013). Even the best contextual stimulus for a neuron typically approximates linear summation of its component inputs(Laboy-Juarez et al., 2019). Although suppression can improve information coding and efficiency(Crochet et al., 2011; Froudarakis et al., 2014; Sachdev et al., 2012; Vinje and Gallant, 2000), sharpen feature selectivity(Angelucci et al., 2017; Jacob et al., 2008) and enhance saliency and boundary detection(Li, 1999a, b), supra-linear summation to highly specific combinations of inputs could represent a powerful means by which cortical neurons might compute and selectively encode higher order features of sensory stimuli. In particular, primary sensory cortical neurons massively outnumber their thalamic inputs, permitting overcomplete representations. Theoretical and experimental work has proposed that overcompleteness permits sparse representations, in which small groups of active neurons efficiently represent natural or complex inputs(Barlow, 1961; de Vries et al., 2020; Lewicki and Sejnowski, 2000; Olshausen and Field, 1997; Stringer et al., 2019). However, how cortical circuits exploit spatial summation to generate these codes is not known. We explored the hypothesis that a population of cortical neurons supra-linearly summating diverse, but specific combinations of inputs could give rise to a sparse, comprehensive code for higher order features, and potentially represent a widespread mechanism in the cortex for the synthesis of complex feature codes. Although much previous work has found that summation across spatially distributed sensory inputs is sub-linear(Angelucci et al., 2017; Armstrong-James and Callahan, 1991; Bair et al., 2003; Blakemore and Tobin, 1972; Boloori and Stanley, 2006; Brumberg et al., 1996; Gilbert, 1977; Higley and Contreras, 2003; Simons, 1985; Simons and Carvell, 1989), by probing large neural populations with spatially diverse stimuli, we find that a majority of cells summate a highly specific subset of stimuli supralinearly, but most other inputs sub-linearly. The diversity and specificity of this supra-linear summation yields a sparse, comprehensive population code for complex spatial features. In particular, each possible combination of whiskers or visual stimulus patches drives activity supra-linearly in a dedicated, similarly sized population of neurons.

## Results

### Stimulus-specific supralinear summation during active sensation in the barrel cortex

We tested the notion of diverse supralinear summation in both the primary somatosensory and visual cortex of the rodent. First we took advantage of the mouse whisker system, which is both discretized and naturally active(Feldmeyer et al., 2013; Petersen, 2007), to probe the logic of cortical integration across stimulus space in awake, active sensing animals. We developed a novel paradigm for reliably generating active touch between a user-defined set of whiskers and a corresponding set of pneumatic actuators (Fig. 1a). Awake mice were head-fixed on a rotary treadmill and habituated to run at relatively high speed (>30 cm/s), a condition in which they moved their whiskers in a rhythmic and stereotyped fashion for many minutes at a time(Pluta et al., 2015; Pluta et al., 2017). All but five of the largest whiskers were trimmed just before the experiment and data was collected in the next 1-2 hours prior to the onset of any trimming induced plasticity(Clem and Barth, 2006; Li et al., 2014). Next, a set of five pneumatically controlled pistons was presented to precise locations, such that on each trial the corresponding set of 1-5 whiskers made repeated active touches with the presented set of pistons (Fig. 1b). On each trial the pistons entered the whisking field within one whisk cycle (~7 ms). Slight trimming of the whiskers ensured that only one whisker made contact with each piston, which we confirmed using high speed imaging (Fig. 1b). An example trial with a video still image and whisker tracking is presented in Figure 1b (and see Extended Data Video 1). In this trial pistons targeted only the B1, C1, and γ whiskers, and each whisker made 17-20 absolutely selective contacts with its respective piston (Fig. 1b). Under these conditions we sampled neural activity in the upper layers with GCaMP6s (Fig. 1c,d see Methods).

**Figure 1.**
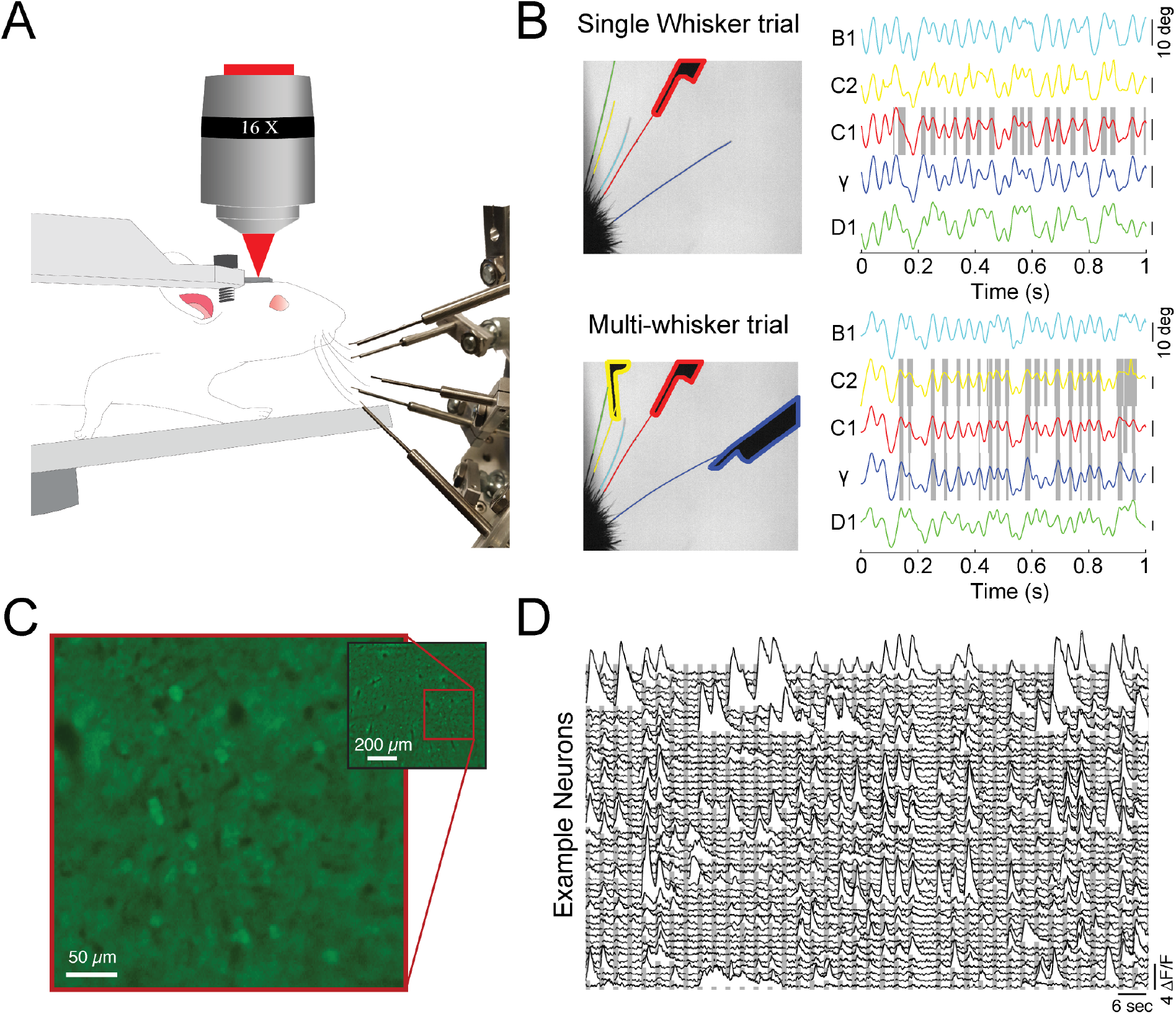
Probing cortical population codes of higher order features during active touch. A) Schematic of the experimental preparation showing a locomoting mouse under a two-photon microscope actively contacting five individual pistons with five individual whiskers (the rest are trimmed just before the experiment, see Methods). B) Top left: single frame of a high-speed video where only the C1 whisker was presented with a piston. Top right: angle traces for each of the five whiskers. Touch events between the C1 whisker and the C1 piston are shaded in grey. Bottom: same as above but for a trial where the C2, C1, and γ pistons were presented. C) Example field of view from a single depth (pixel values are the variance across time). D) Two minutes of consecutive fluorescence measurements from 50 randomly chosen neurons in a single example experiment. Vertical gray bars indicate presentation of different multi-piston stimuli.

To probe how barrel cortex neurons encoded first and higher order tactile features we presented all 31 possible combinations of the five pistons. 48% (2262/4756) of identified neurons showed a significant response (see Methods) to at least one of the 31 stimuli. When the pistons were presented individually to map the ‘receptive field’ of each touch responsive neuron, 52.7% of neurons exhibited a significant response to just one of the single whisker stimuli, 12.6% responded to 2 or more, and a striking 34.8% did not respond to any single whisker stimulus, but responded robustly to at least one multi-whisker combination. Both L4 and L2/3 excitatory neurons organized into loose spatial maps of preferred whiskers consistent with prior findings from anesthetized mice(Clancy et al., 2015; Kerr et al., 2007; Sato et al., 2007) (Fig 2a and Fig. S1).

**Figure 2.**
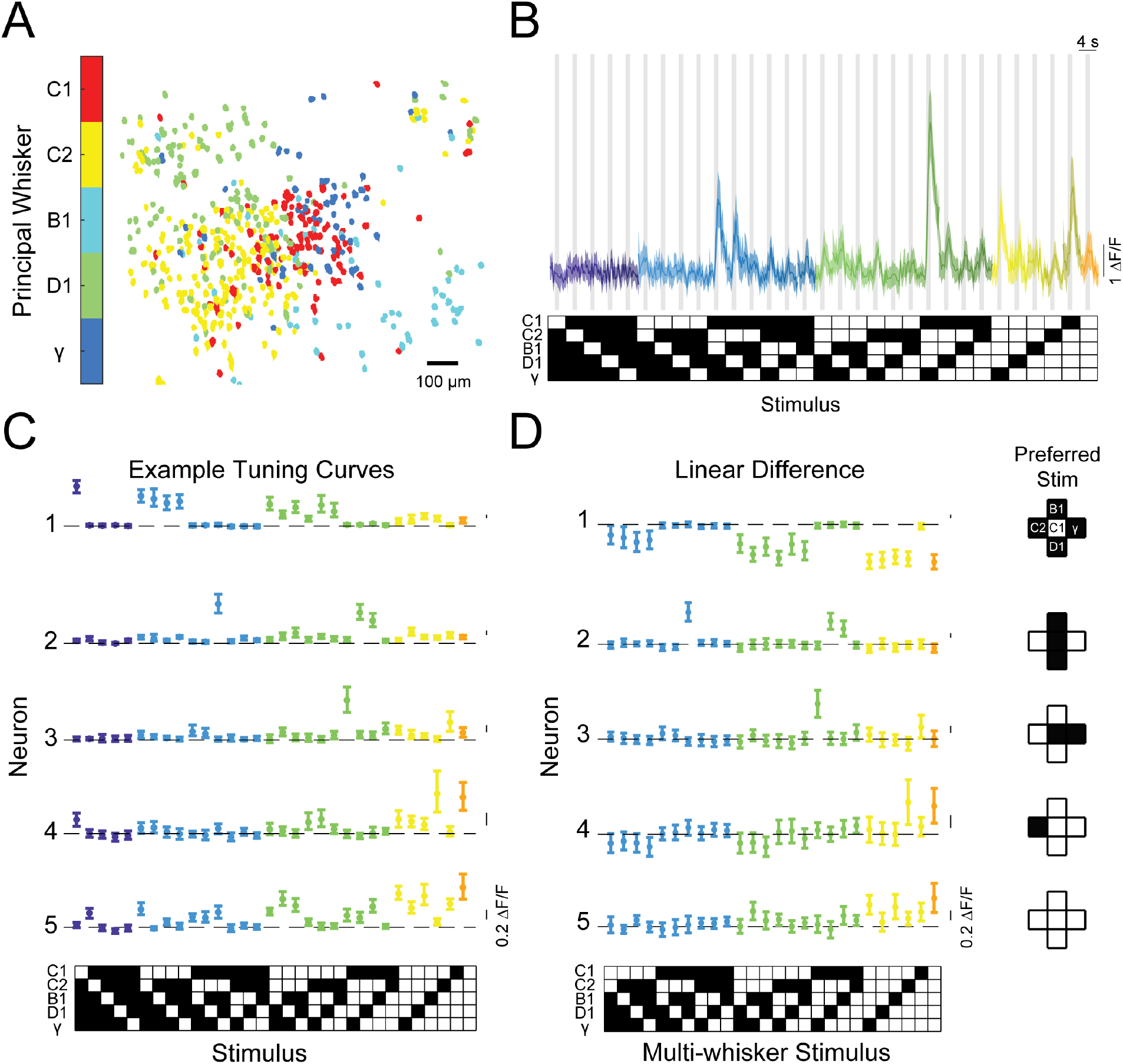
Diverse and selective representations of higher order tactile stimuli in mouse barrel cortex. A) Example map of principal whisker preference in L2/3 of one mouse. B) Top: Calcium responses from an example neuron to each of the 31 possible combinations of the five pistons (mean and 95% C.I.). Traces are colored-coded by the number of whiskers contacting pistons in each stimulus (1-5 whiskers). Vertical gray bars indicate presence of the tactile stimulus. Bottom: Diagram of the 31 corresponding piston combination stimuli. White squares indicate the presence of the corresponding piston and black indicates the absence. C) Example tuning curves for 5 neurons in L2/3 of S1 for all 31 stimuli and colored as in B). Neuron 3 corresponds to the neuron shown in panel B). D) Left: The computed difference of the observed response to each multi-whisker stimulus from the linear sum of the component responses when each whisker contacts its corresponding piston alone for the five example neurons in C), or ‘Linear Difference’. All error bars are bootstrapped 95% confidence intervals (C.I.). Right: Diagram of each neuron’s preferred stimulus.

In many previous recordings from the barrel cortex in which whiskers were passively stimulated, stimuli presented outside a neuron’s receptive field typically suppressed the response to stimulation of the receptive field alone(Arabzadeh et al., 2003; Brumberg et al., 1996; Drew and Feldman, 2007; Fanselow and Nicolelis, 1999; Laboy-Juarez et al., 2019; Sachdev et al., 2012) (but see others(Estebanez et al., 2016; Ramirez et al., 2014; Shimegi et al., 1999)). We observed a strikingly different type of modulation in most responsive neurons. Consider the neuron whose calcium responses are shown in Fig. 2b. This neuron showed no significant response to any single piston alone, but exhibited the strongest activity when pistons targeting the C2, B1, D1 whiskers were presented in combination. Tuning curves for five example neurons are shown in Fig. 2c. The first neuron showed a conventional tuning curve, exhibiting a strong and selective response to the C1 piston when presented alone, but weaker responses when any other pistons were presented in combination with the C1 piston. In contrast, neurons 2-5 (neuron 3 is the same as in Fig. 2b) all showed preferred responses for different multi-piston combinations. Remarkably, for these other neurons most other combinations of pistons had a weak or suppressive effect on the neuron’s activity, giving rise to a highly selective response for their specific multi-whisker stimulus.

We next computed the difference between the actual response of each neuron to each multi-whisker stimulus, and the predicted response based on a linear sum of the responses to individual whisker stimuli (‘Linear difference’, Fig. 2d). For all five neurons, most multi-whisker stimuli drove sub-linear summation (i.e., suppression) or had no effect. However, for the last four of these five example cells, the stimulus that drove the largest response across the full 31 stimulus set showed a pronounced supra-linear effect, in that the pistons in each preferred combination drove much larger responses than the linear sum of their component pistons when presented alone.

### A comprehensive population code for higher order tactile stimuli in the upper layers of the barrel cortex

If many other neurons were to show similarly strong and selective responses for various combinations of whisker touches, with different neurons encoding all possible combinations, barrel cortex could contain a sparse, comprehensive code for higher order tactile features in the stimulus space we probed (what has been considered an ‘overcomplete’ code since the number of responsive neurons in S1 exceeds the stimulus dimensionality(Olshausen and Field, 1997)). To test this possibility, we asked whether these large populations of barrel cortex neurons encoded all 31 possible piston combinations within the five-piston stimulus set (5 single whisker and 26 multi-whisker stimuli), thus covering the stimulus space, or if instead specific combinations were strongly over-represented. Fig. 3a shows tuning curves of 31 neurons from one mouse that covered the stimulus space. Importantly, across all imaged mice, all 31 stimuli were represented approximately evenly (Fig. 3c). To ensure the robustness of this analysis, we computed each neuron’s preferred stimulus on 50% of the data and used it to order tuning matrices computed on the remaining 50% (see Methods). This demonstrates that barrel cortex has the ability to sparsely encode all possible higher order tactile stimuli in the five-whisker space we probed. Comparing the stimulus selectivity and deviation from linear summation from each neuron side by side shows that the population code for multi-whisker stimuli correlated well with selective supra-linear summation for the most preferred stimulus (Fig. 3b,d). This supports the idea that stimulus-specific summation over surround whiskers helps promote a sparse and selective code for higher order tactile features: for most whisker combinations summation was sub-linear (i.e., suppressive, consistent with prior findings). However, for a small subset of multi-whisker inputs summation was strongly supra-linear. Another important rule emerged from this analysis: the best stimulus for each neuron nearly always contained the principal whisker (PW), demonstrating that it was required for the preferred supra-linear response (71% of neurons in C1 column, see Methods, Fig. S2a,c). The preferred stimulus also almost always included the anatomically aligned ‘columnar’ whisker (CW) even when the CW and PW were different (60% of C1 neurons’ PW was C1, Fig. S2b).

**Figure 3.**
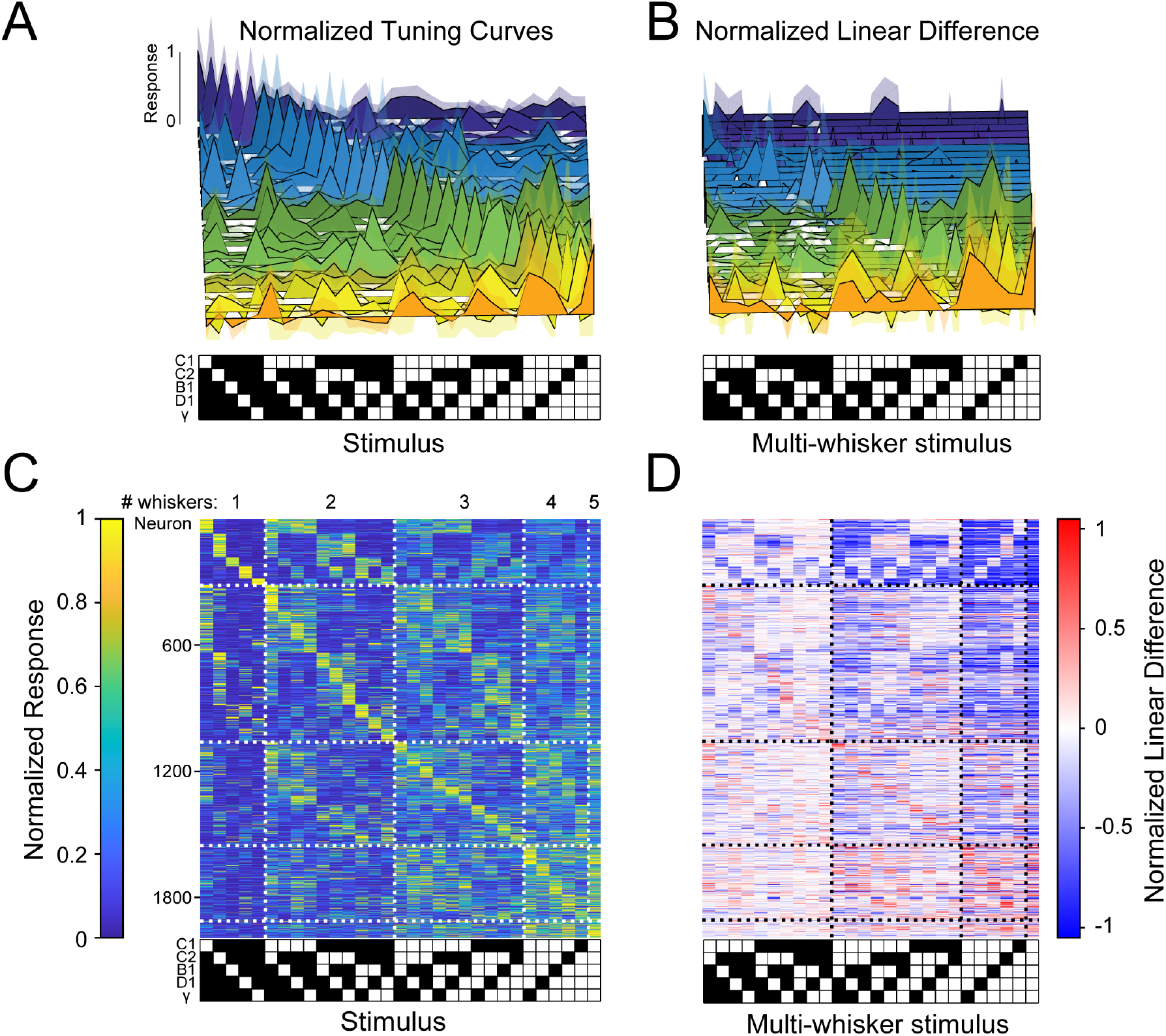
Sparse, high dimensional population codes of higher order tactile features generated through stimulus-specific supra-linear summation. A) 31 example tuning curves from one field of view in one mouse showing neurons that exhibit a peak response for each of the 31 presented stimuli. Shaded regions are 95% C.I. B) Linear difference of the 31 neurons in A). C) Normalized, cross-validated tuning curves for all sensory-driven neurons in awake, whisking mice. Neurons are ordered along the vertical axis by the stimulus to which they responded best. D) The normalized linear difference for the neurons in C).

We next compared the population codes in L4 (2,428 neurons) and L2/3 (2,328 neurons): first we found neural ensembles in both layers encoded the complete set of the 31 presented second order tactile stimuli (Fig. S3a,b), implying that even at the first stage of cortical processing cortical integration gives rise to a sparse, comprehensive code of higher order stimuli. However, we observed several important differences indicative of a translaminar transformation of the neural codes of these features. In L4, the principal whisker contributed substantially more to the individual neurons’ responses than in L2/3, and explained much more of the variance in their responses to the stimulus set. Conversely, in L2/3, the surround whiskers, often in highly specific and non-contiguous combinations, shaped the tuning to multi-whisker stimuli. Ultimately, as a consequence, a much greater fraction of L2/3 neurons responded more specifically to unique combinations of whiskers, than in L4. More than three times as many L2/3 neurons as in L4 exhibited multi-whisker receptive fields (Fig. S3c), and L2/3 neurons were substantially less selective for single whisker stimuli (Fig. S3d,e). Strikingly, L2/3 had nearly three times the number of neurons that exhibited driven responses for a multi-whisker stimulus that did not include the PW (14% vs. 43%; Extended Data. Fig. S3f,g) implying a more diverse code for complex tactile features. L2/3 also had a greater fraction of neurons whose preferred stimulus lacked the ‘columnar whisker’ (analysis restricted to the C1 column, Fig. S3h). Similarly, among these C1 column neurons, the contribution of the C1 whisker to neuronal tuning curves was significantly more distributed than for L4 neurons (Fig. S3i). These differences suggest that L2/3 could better represent the combinations of whiskers in a given stimulus with a more spatially compact population of neurons.

### A similar mechanism also generates a comprehensive population code for higher order stimuli in the mouse visual cortex

If the synthesis of higher-order feature codes through stimulus-specific spatial summation is a widespread feature of cortical population activity, then we should be able to observe a similar phenomenon in a cortical area that receives extremely different sensory input, such as the primary visual cortex. To test this hypothesis, we made recordings of neural activity in L4 and L2/3 of mouse V1 while presenting five patches of drifting gratings, each approximately the size of the average classical receptive fields of most mouse V1 neurons, 10 visual degrees (see Fig. 4a and Methods). The drift direction of the center patch was orthogonal to those in the surround patches to ensure robust visual responses (Cavanaugh et al., 2002; Self et al., 2014; Shushruth et al., 2013). As in barrel cortex, responses to single patch stimuli reflected the underlying topographic map (Fig. 4b). In agreement with the notion of diverse, stimulus-specific summation, we likewise observed that V1 populations encoded all possible combinations of the five stimulus patches in approximately even fashion (Fig. 4c,e). Moreover, just as in the barrel cortex, many V1 neurons summed these grating patches supra-linearly to give rise to their preferred stimulus, but summed nearly all other stimuli sub-linearly, effectively sharpening the tuning to the preferred stimulus (Fig. 4d,f). As in barrel cortex (Fig. S4), this coding principle was shared across layer 4 (Fig. S5a,b) and layer 2/3 (Fig. S5c,d). To ensure that this was not specific to the precise patch size of 10 degrees, we also tested 15 degree patches, and observed qualitatively similar results (Fig. S6).

**Figure 4.**
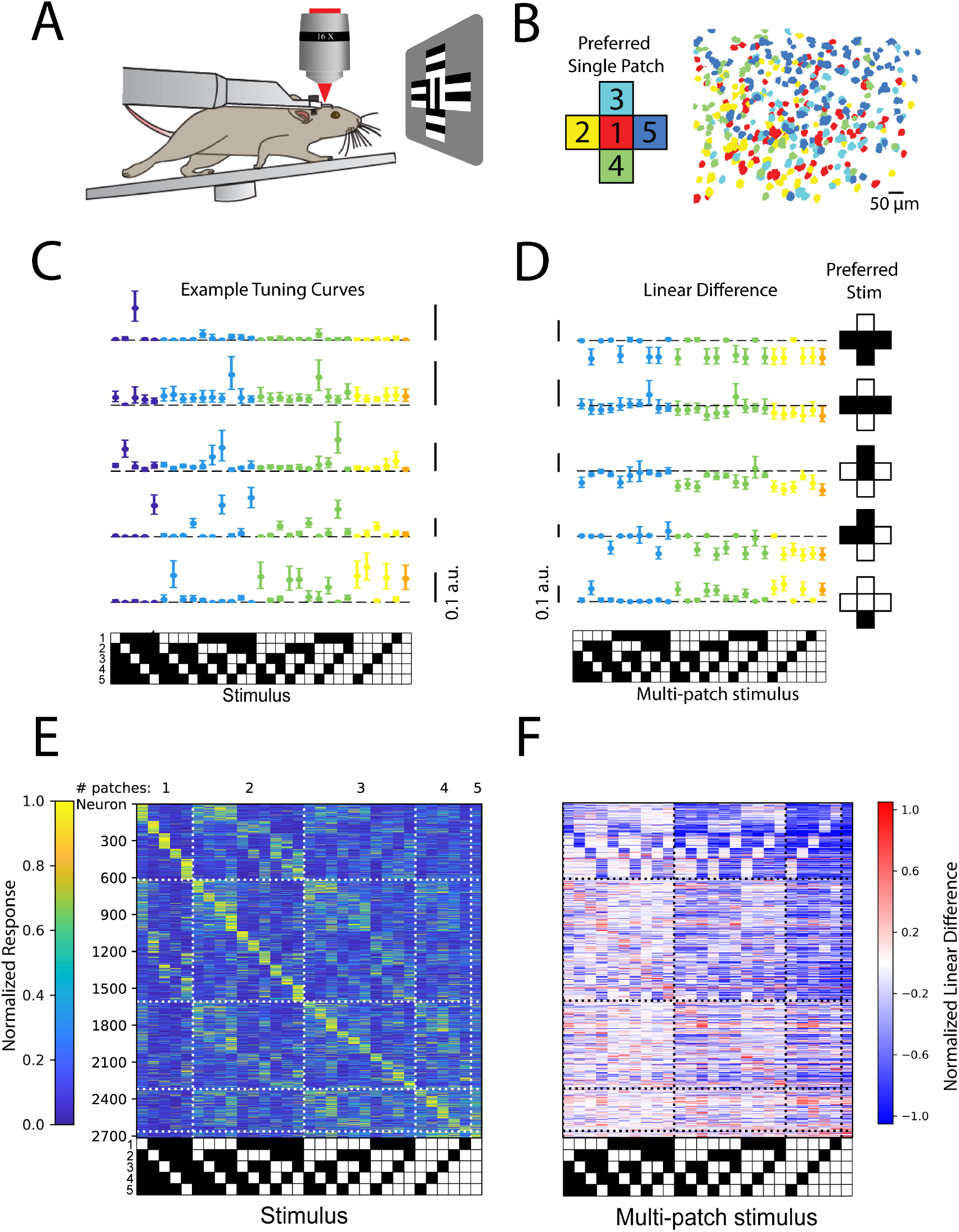
Stimulus-specific supra-linear summation in the primary visual cortex. A) Schematic of the experimental preparation. B) Example imaged field of view in layer 2/3 where each identified neuron is color coded according to its preferred single stimulus patch. C) Five example tuning curves from neurons preferring different numbers of patches. Neurons are from four fields of view in layer 2/3 of three mice. Error bars are bootstrapped 95% C.I. D) Linear difference of the five neurons in C). E) Normalized, cross-validated tuning curves for all sensory-driven neurons in awake, whisking mice. Neurons are ordered along the vertical axis by the stimulus to which they responded best. F) The normalized linear difference for the neurons in E).

In order to quantify the degree to which population codes reflected supra-linear vs. sub-linear summation, we first sorted the stimuli in order of preference for each neuron (Fig. 5a). Then, we computed linear differences as before for multi-whisker or multi-patch stimuli (Fig. 5b). By averaging the linear difference of ranked stimuli across neurons, we were able to quantify how supra- or sub-linearity of multi-whisker or multi-patch stimulus response varied with stimulus preference. We found that in multi-whisker or multi-patch preferring neurons across both cortical areas, sub-linear summation dominated for all but the most preferred stimuli (Fig. 5c,d). In contrast, supra-linear summation was highly selective for only the most preferred stimulus on average across both cortical areas and layers.

**Figure 5.**
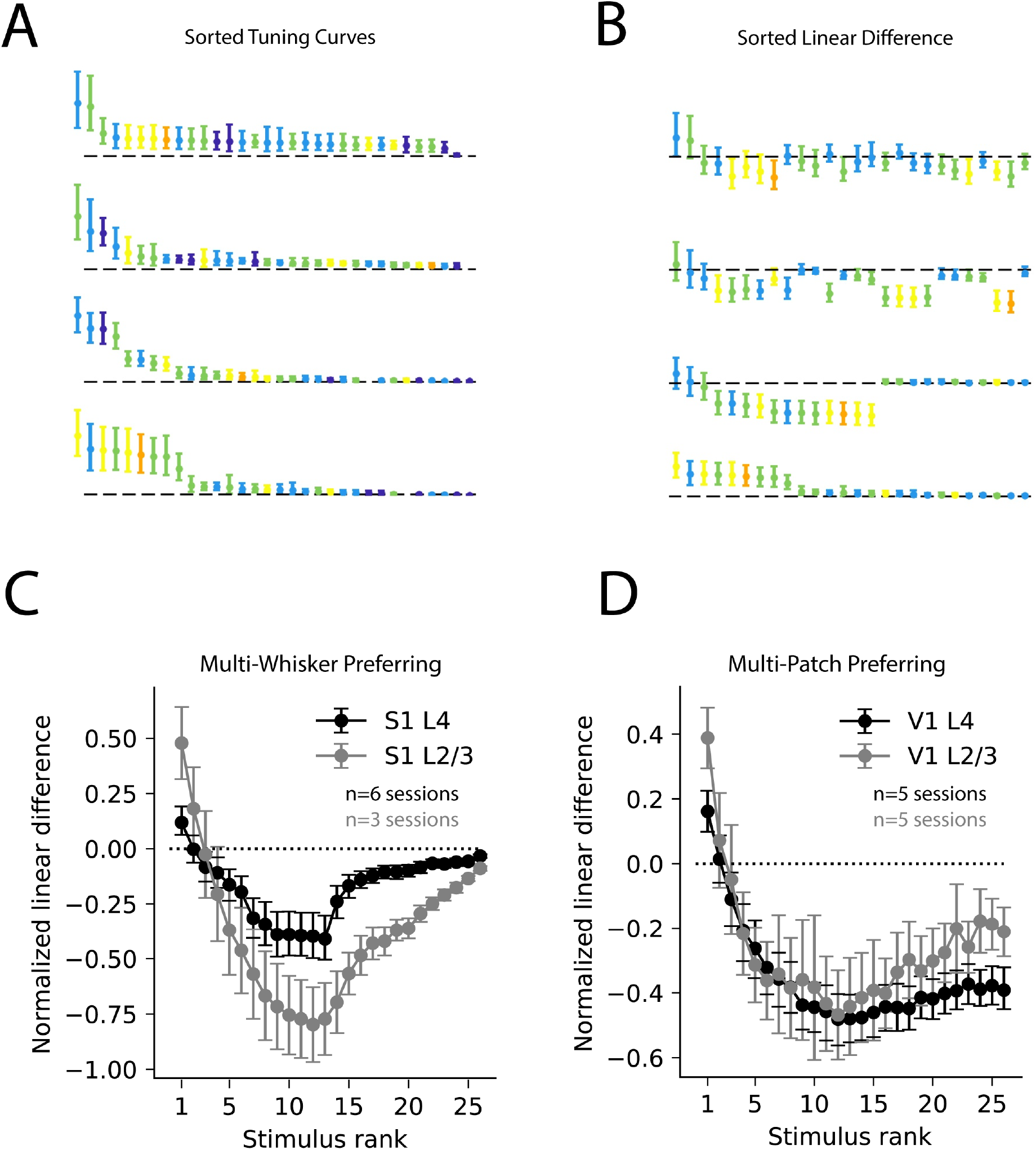
Selective superlinearity and global sublinearity of spatial summation across cortical areas and layers. A) Example V1 layer 2/3 tuning curves of the four multi-patch preferring neurons reproduced from Figure 4C, sorted by descending response magnitude. B) Linear difference of the same neurons’ multi-patch stimulus responses, sorted as in A). C) Sorted linear difference averaged across neurons that preferred a multi-whisker stimulus in S1 L4 and L2/3. D) Same as C) for V1 L4 and L2/3.

To ensure that our estimation of neuronal responses from calcium signals to the stimulus set is not substantially compromised by significant non-linearities in GCaMP6s, we calibrated the transform from action potential rate to GCaMP6s fluorescence changes using two-photon targeted loose patch electrophysiology in CamK2a-tTa;tetO-GCaMP6s mice (see Methods, experiments were conducted in the visual cortex for technical reasons). We found this transform to be approximately linear and highly sensitive to firing rate changes (mean Pearson’s correlation coefficient = 0.71 ± 0.03, mean ± s.e.m., n=21), consistent with prior findings(Chen et al., 2013) (Fig. S7). For our barrel cortex data, we also tracked all five whiskers, including touches and bend, for all trials in the dataset, using an artificial neural network approach (DeepLabCut(Mathis et al., 2018), see Methods) to determine if sensorimotor adaptation in how the whiskers contacted the pistons could have explained the non-linear summation we observed in our neuronal responses.

In a subset of mice (n = 3) we measured five key parameters of the center whisker (C1): mean number of touches per trial, mean angle during the contact period, mean whisker bend, the amplitude of whisking, and the mean duration of contact. First, we compared these five kinematic variables between trials when individual pistons were presented alone with trials when each of the 26 combination stimuli were presented. However, we observed no difference in the number of touches, mean bend, or whisk amplitude, but did detect a decrease in mean contact duration and mean angle with increasing number of pistons (see Fig. S8 and Methods). These data imply that stimulus-specific behavioral adaptation in the mouse’s whisking could not have contributed to the stimulus-specific supra-linear facilitation we observed, although it may have led to a general trend of modestly weaker sensory responses for stimuli containing an increasing number of pistons that would be systematic across all neurons.

Brain state and behavioral state can profoundly influence cortical computation. Whether they also effect how cortical neurons integrate over space to encode higher order stimuli remains uncertain. In the barrel cortex, we compared spatial summation over the whisker array between the awake, active whisking state described above with passively stimulated conditions in anesthetized mice. In the barrel cortex of lightly anesthetized mice (to prevent active whisking) we stimulated the same five whiskers with a set of five piezo actuators (see Methods). Although multiple previous studies have used noise stimuli(Brumberg et al., 1996; Estebanez et al., 2012; Ramirez et al., 2014) or specific deflections (Higley and Contreras, 2005; Laboy-Juarez et al., 2019; Mirabella et al., 2001b) in anesthetized or sedated animals to estimate neuronal receptive fields and the impact of surround whiskers in S1, we used mechanical stimuli that approximated the movements of the whiskers during active sensation to more specifically compare barrel cortex coding between active and passives states. Population responses in these anesthetized mice were not as uniformly distributed across multi-whisker space as in awake, whisking mice, and instead favored broader spatial summation, or in other words, exhibited poorer feature selectivity (Fig. S9a,c,e). Furthermore, L2/3 tuning curves in awake mice were substantially less well fit by a general linear model, potentially owing to enhanced nonlinearity in the awake, active condition (Fig. S9f). These results imply that anesthesia and passive touch linearize summation in L2/3 of the barrel cortex and alter the population code of higher order features. Taken together, our data demonstrate that brain and behavioral state potently modulate how somatosensory populations respond to higher order stimuli.

## Discussion

Our data show that the logic of spatial summation in the sensory cortex leads to the generation of a distributed feature code of more complex tactile or visual stimuli. Unlike most prior studies, we found that the firing rate of barrel or visual cortical neurons, particularly in the awake, active state, could be potently facilitated by surround stimuli, and many neurons could only be driven by more complex features. Importantly, this surround facilitation was highly specific to just one or a small number of multi-whisker combinations, while most other combinations suppressed or had no impact on the response to preferred stimuli. The net consequence was to generate highly selective responses for specific tactile or visual ‘features’ composed of active touches with a specific set of surround whiskers or grating patches that extended well beyond the classical receptive field. Furthermore, a small, but relatively uniform fraction of the imaged neurons encoded each of the presented stimuli, and as a population these neurons encoded a sparse, but comprehensive representation of the stimulus space.

This potent facilitation by stimulation of specific components of the receptive field surround contrasts with the widely observed phenomenon of surround suppression, a prominent form of divisive normalization(Carandini and Heeger, 2013). In our data, surround suppression captures the response of many neurons to their non-preferred multi-whisker stimuli, but not for their preferred stimulus. Taken together, our data imply that sparse, distributed ensembles of barrel or visual cortical neurons might encode a core set of higher order features in whisker or retinotopic space from which more complicated representations could be built, perhaps in higher cortical areas. These ensembles in the primary sensory cortex might therefore be crucial for contour detection and shape perception of three-dimensional objects.

Our findings also raise the possibility that stimulus-specific supra-linear summation might be a more general feature of cortical sensory coding, or even cortical coding more generally, in active contexts. Future work, perhaps combining functional connectomics(Bock et al., 2011) and multiphoton optogenetics(Mardinly et al., 2018; Papagiakoumou et al., 2010), could elucidate the precise circuit mechanisms that give rise to such stimulus specific supra-linear summation. In the mouse visual cortex, neurons with preferences for similar visual stimuli show a significantly greater tendency to synaptically couple(Ko et al., 2011), which may lead to recurrent amplification of feature selectivity. Perhaps even more specific excitatory subnetworks could explain the generation of the higher order feature codes observed here, potentially relying on the target specificity of horizontal projections(Gilbert and Wiesel, 1983; Li and Gilbert, 2002; Petersen and Crochet, 2013). Additionally, stimulus-specific modulations in sensory adaptation might contribute to the generation of feature selectivity, as is true in paralyzed rats, although this was only reported for neurons in deeper layers(Ramirez et al., 2014). Finally, feedback from higher cortical areas or secondary thalamic nuclei(Jouhanneau et al., 2014; Williams and Holtmaat, 2019), perhaps conveying signals related to self-motion or sensory predictions, might be critical for generating the supra-linear response to specific complex stimuli, perhaps through non-linear activation of pyramidal cell dendrites(Petreanu et al., 2012; Xu et al., 2012). Such dendritic activity might be facilitated by local dendritic clustering of specific afferents(Iacaruso et al., 2017; Takahashi et al., 2012; Wilson et al., 2016), or circuits for dendritic disinhibition that are recruited by top-down motor-sensory projections(Lee et al., 2013). In addition to understanding its detailed circuit mechanisms, future work can address how higher order feature selectivity in the primary sensory cortex gives rise to ever more selective representations for the identification of specific objects.

---

**Fig. S1.**
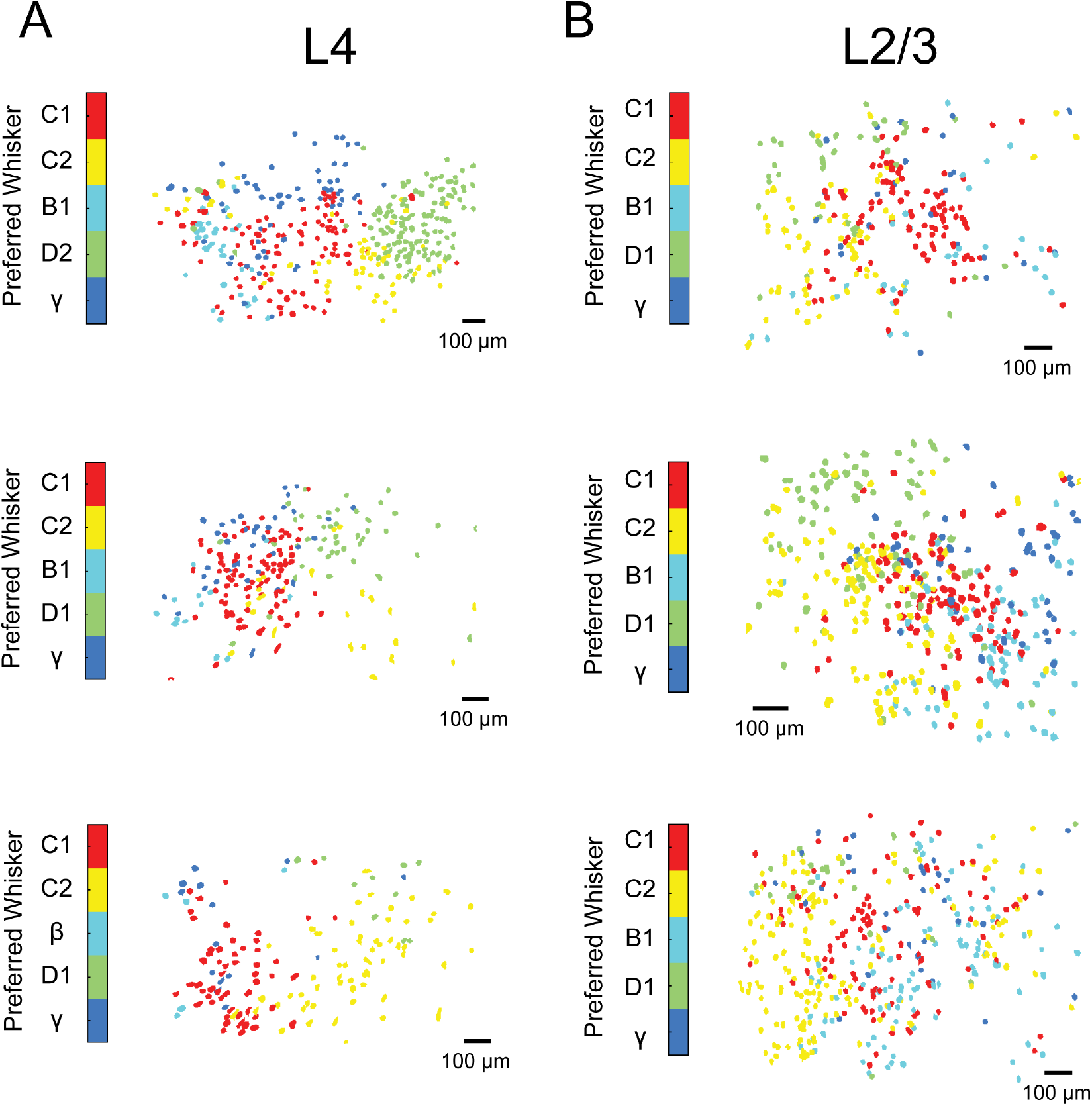
Maps of single whisker tuning in awake, whisking mice. A) Three maps of principal whisker preference for L4 neurons from three Scnn1-Cre mice injected with AAV-syn-flexed-GCaMP6s. B). Same as a) but for three mice recorded in L2/3 using Camk2a-tTA;tetO-GCaMP6s mice.

**Fig. S2.**
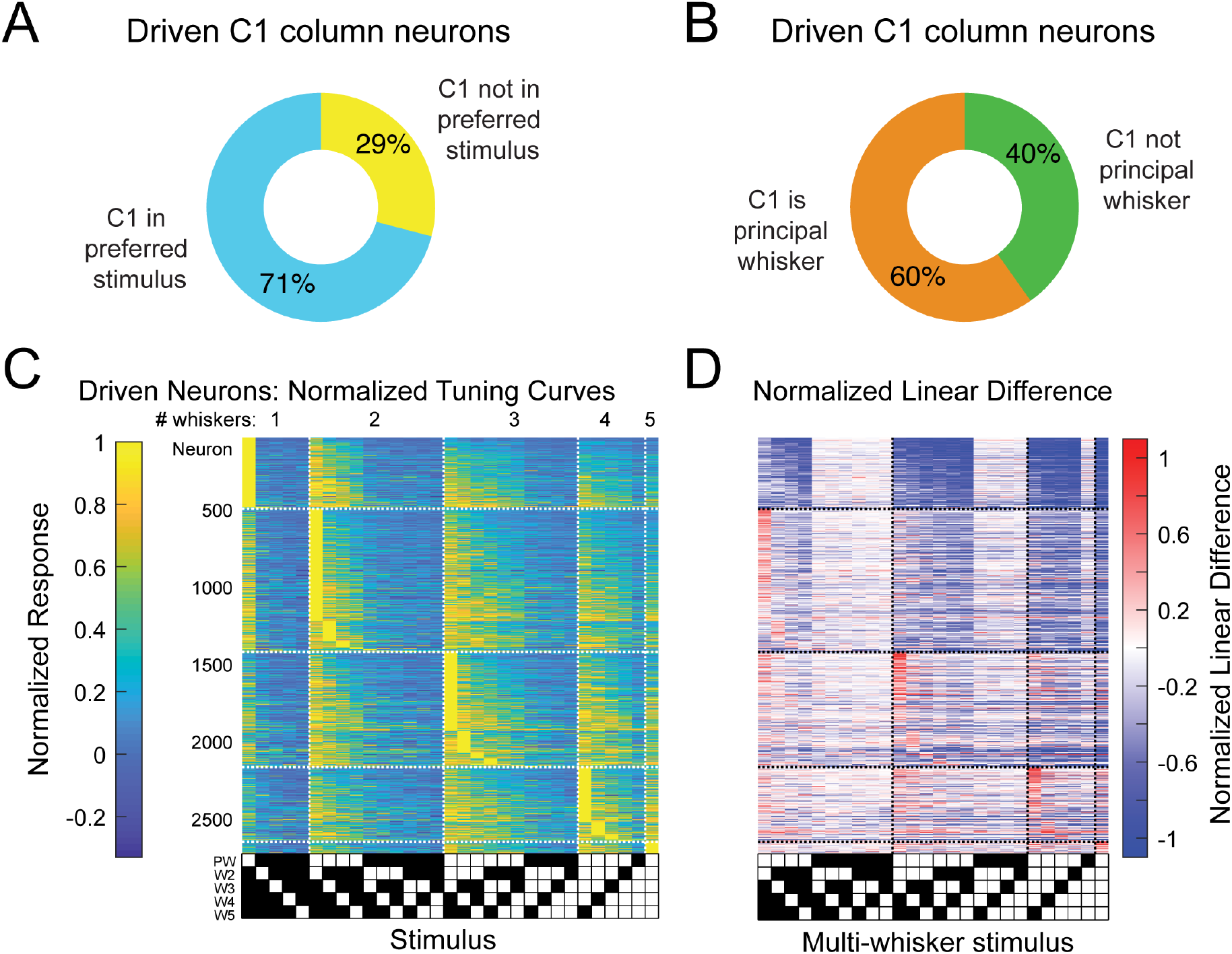
The preferred multi-whisker stimulus for each neuron nearly always includes its principal whisker. A) Breakdown of C1 column neurons by whether they have C1 in their best stimulus or not. B) Breakdown of C1 column neurons by whether C1 is their principal whisker. C) Normalized tuning curves for all sensory-driven neurons in awake, whisking mice. Neurons are ordered on the vertical axis first by the number of whiskers in their preferred stimulus (1-5), and secondly by their selectivity. The principal whisker (PW) is set as the whisker that has the largest weight when fit by a general linear model. The stimuli are ordered by the whisker rank of each neuron’s GLM fit (which is specific to each neuron). Note that the maximal values for each neuron’s tuning curve (rows of the matrix) almost always contains the PW. D) Normalized linear difference for the neurons in A) ordered in the same way.

**Fig S3.**
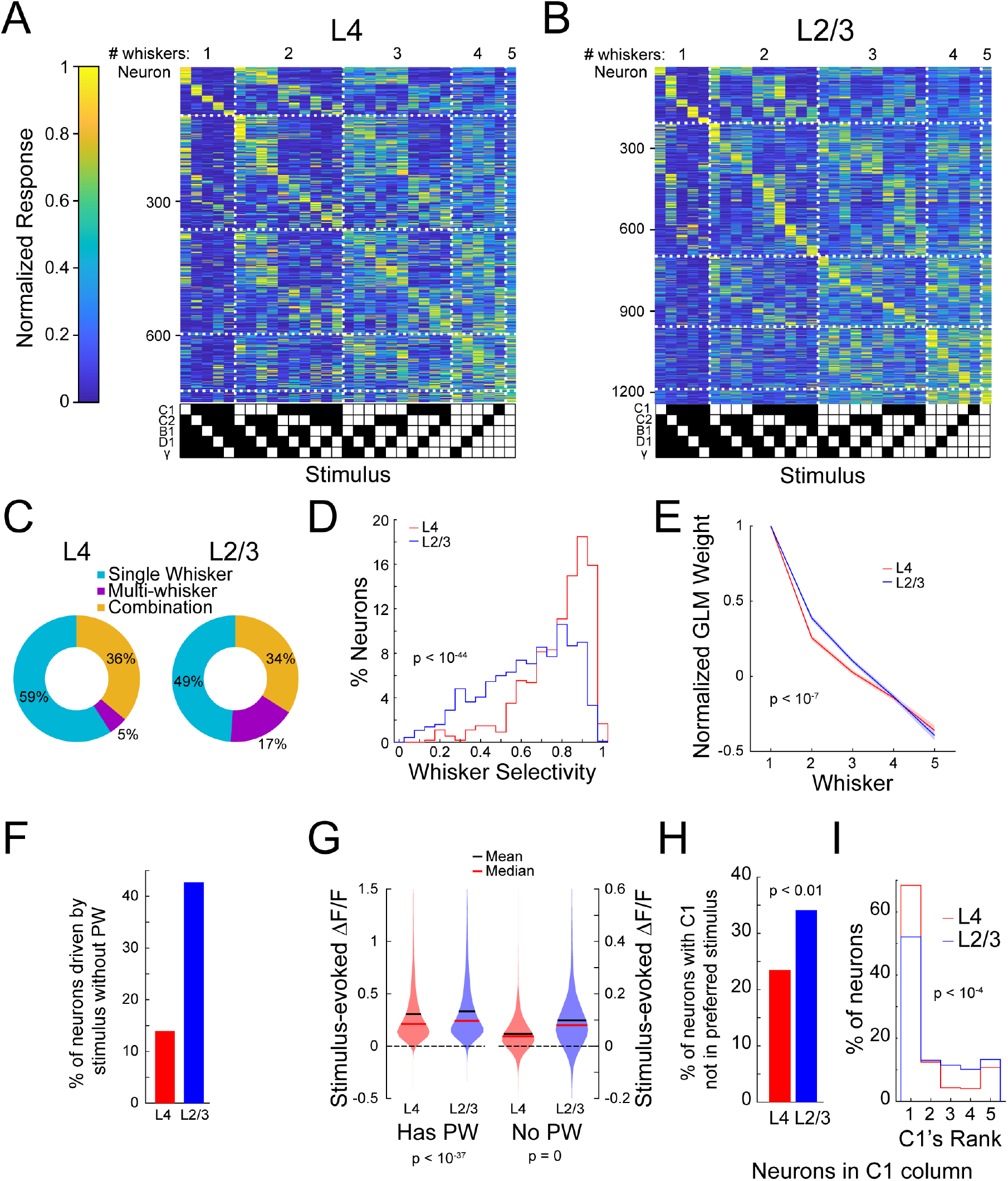
Differences between tactile representations in L4 and L2/3. A) Normalized, cross-validated tuning curves for recorded L4 neurons. B) Same as A) but for L2/3 neurons. C) Ring graphs showing the percentage of driven neurons that were single-whisker responsive, responded to multiple single whisker stimuli, or did not respond to any single whisker stimulus but responded to a multi-whisker stimulus (‘combination’). D) Histograms comparing the whisker selectivity of L4 and L2/3 neurons that are significantly driven by at least one single-whisker stimulus (p < 10^−44^, Wilcoxon ranksum; L4 n=541, L2/3 n=935). E) Average of normalized, rank-ordered whisker weights from general linear models fit to each neuron (p < 10^−7^, 2-way ANOVA; L4 n=847, L2/3 n=1415). F) Percentage of neurons that had at least one significant response to a stimulus that did not include the PW. G) Left: Violin plots of the mean responses for all multi-whisker stimulus combinations for all driven neurons containing the PW (p < 10^−37^, Wilcoxon ranksum; L4: n=12705, L2/3: n=21225). Right: Same but for stimuli that did not contain the PW (p = 0, Wilcoxon ranksum; L4: n=9317, L2/3: n=15565). H) Percentage of C1 column neurons that did not have C1 in their preferred stimulus (p < 0.01, permutation test; L4 n=345, L2/3 n=384). I) Distribution of C1 column neurons’ rank of the C1 whisker within their GLM-computed whisker weights (p < 10^−4^, Kruskal-Wallis test; L4 n=345, L2/3 n=384).

**Fig. S4.**
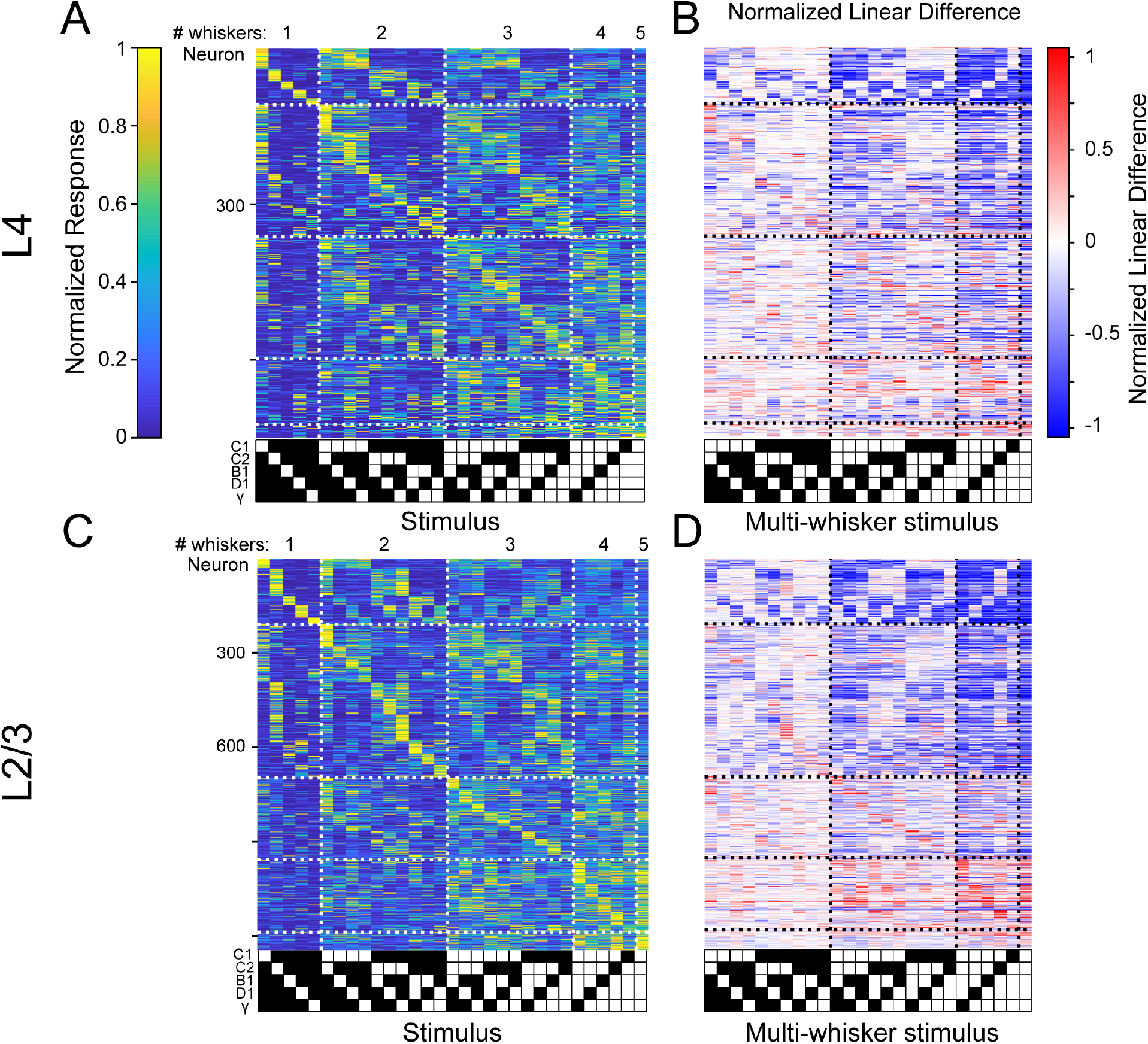
Comparison between L4 and L2/3 of barrel cortex. A) Normalized tuning curves of 50% of the data for all sensory-driven L4 neurons from awake mice ordered on tuning curves calculated on the remaining 50% of data. The black diagonal line depicts the expected population diagonal if the neural population uniformly covers the stimulus space. B) Normalized linear difference for the neurons and data shown in A). C) Same as A) but for L2/3 neurons recorded in awake mice. D). Normalized linear difference for the neurons in C).

**Fig. S5.**
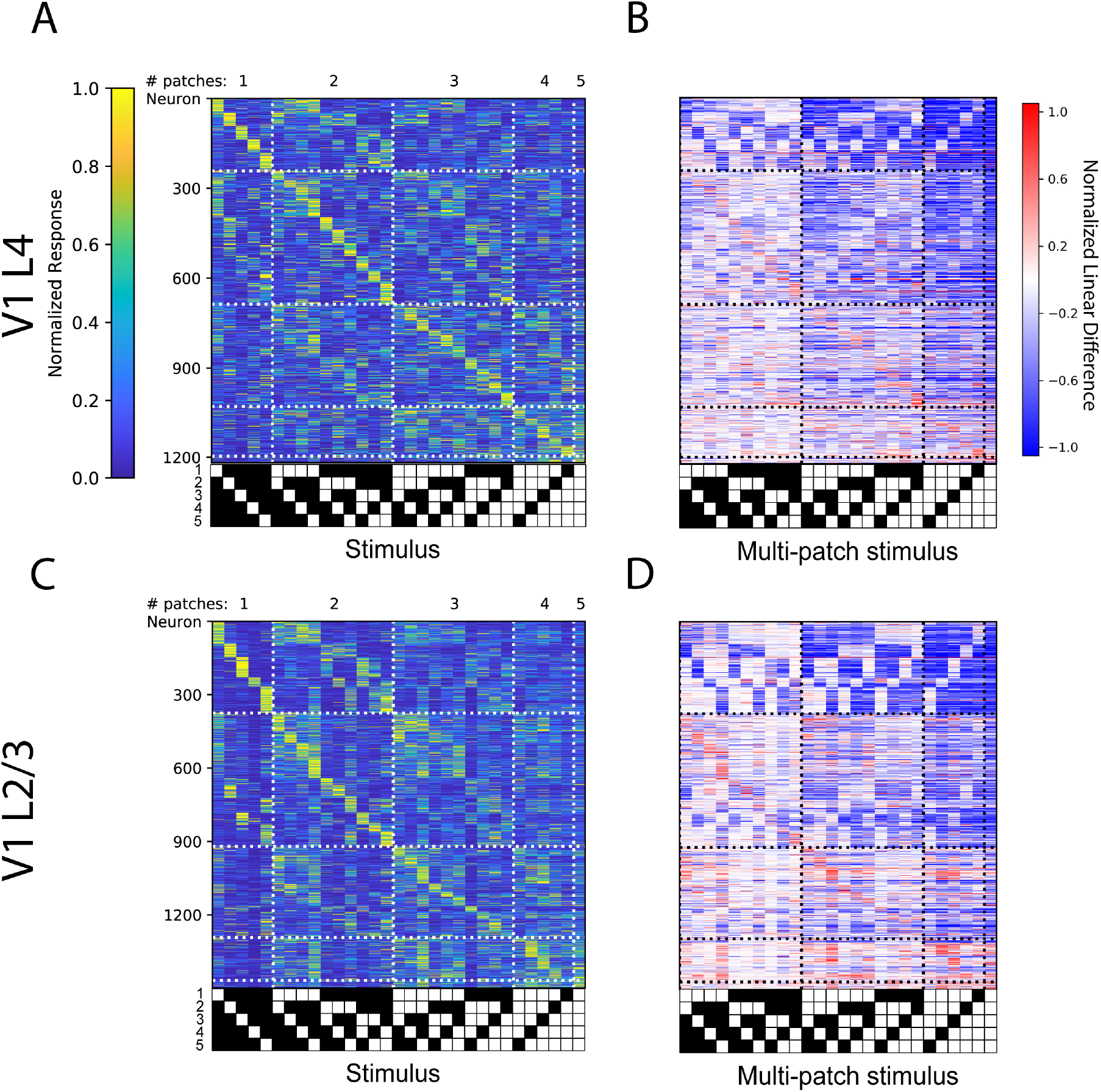
Comparison between L4 and L2/3 of visual cortex. A) Normalized tuning curves computed from 50% of the data for all sensory-driven V1 L4 neurons from awake mice ordered on tuning curves calculated on the remaining 50% of data. B) Normalized linear difference for the neurons in A). C) Same as A) but for V1 L2/3 neurons recorded in awake mice. D). Normalized linear difference for the neurons in C).

**Fig. S6.**
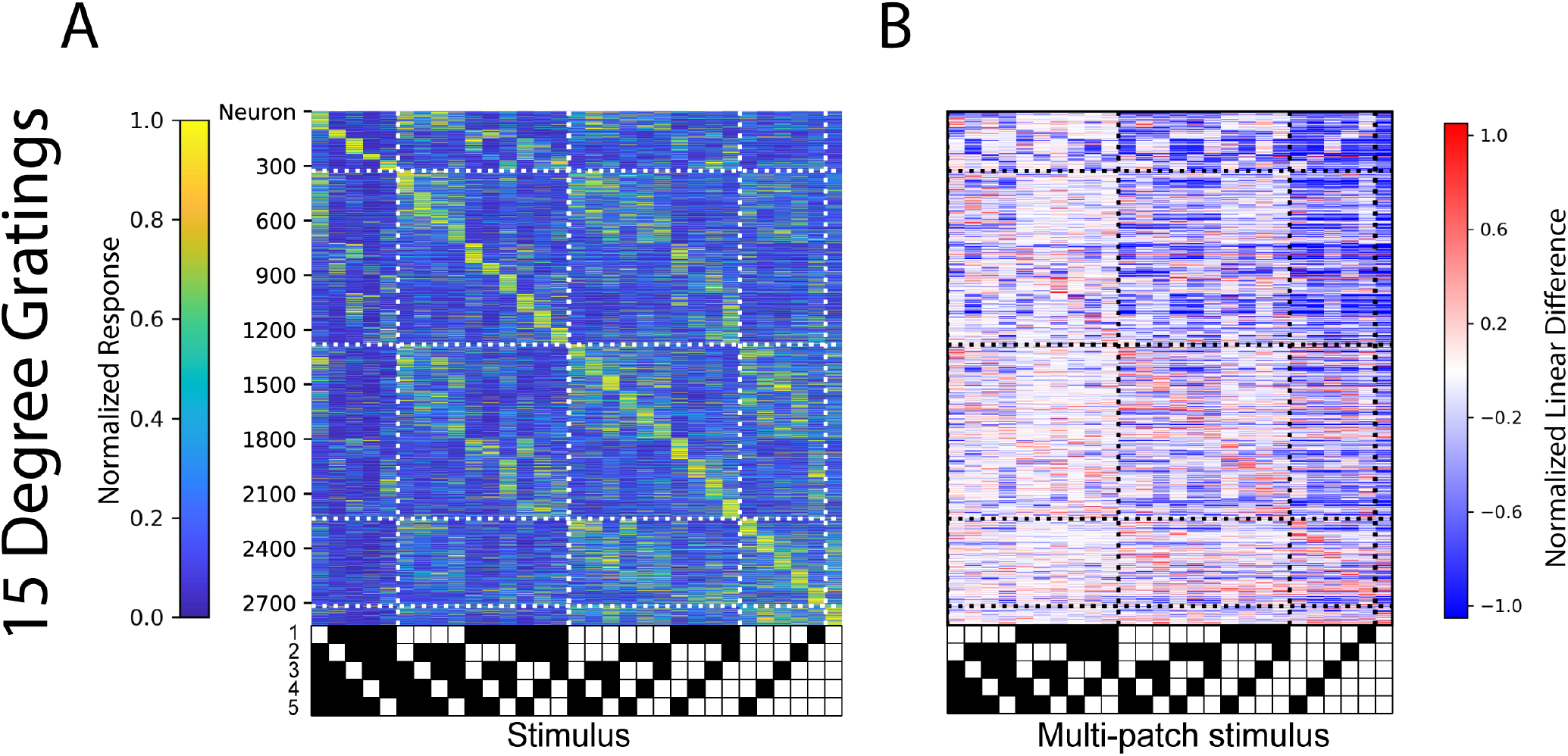
Analogous population code for 15 degree patches. **A)** Cross-validated tuning curves and B) linear difference plots of neurons in layer 4 and layer 2/3 of V1 for 15 visual degree patches (right).

**Fig. S7.**
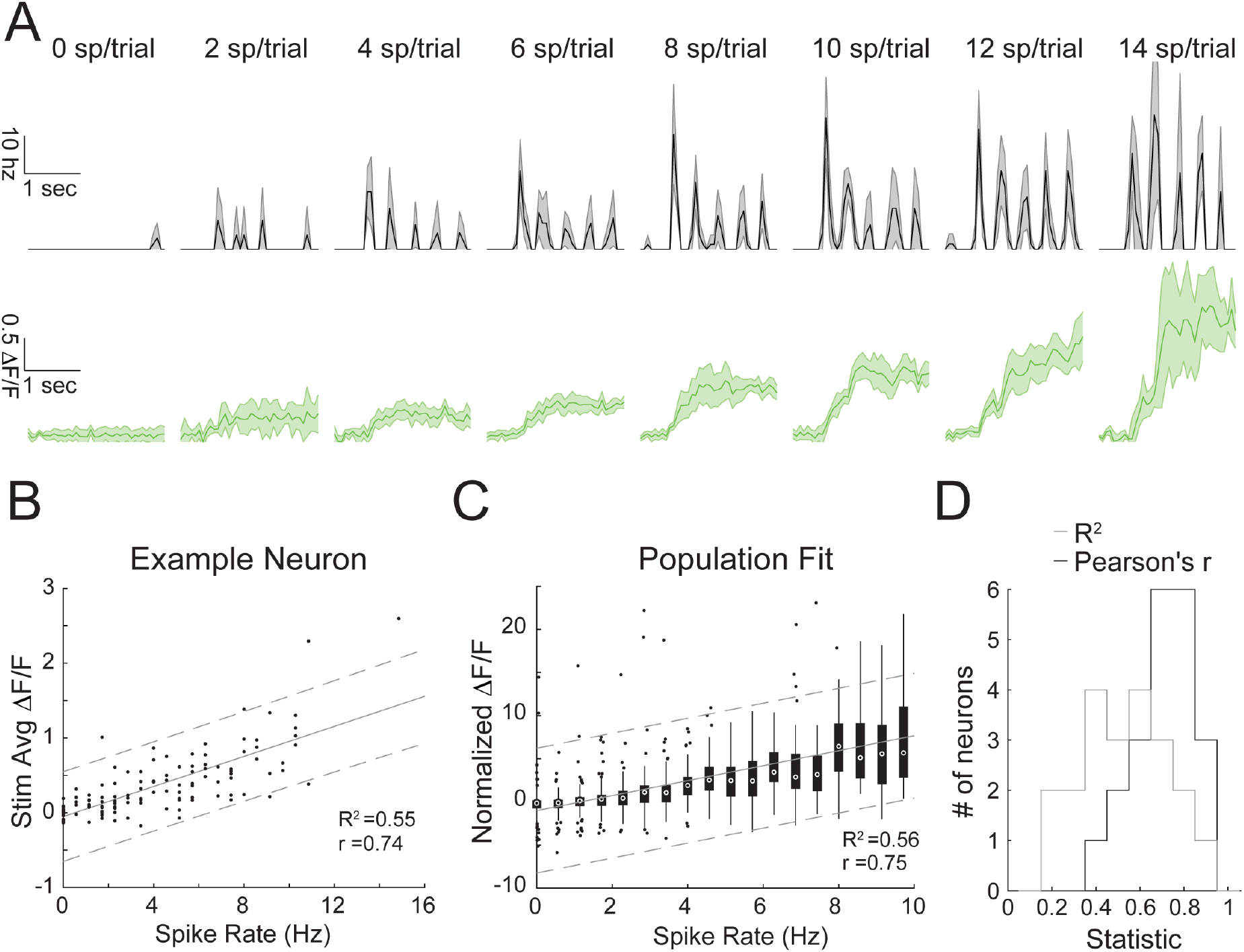
The transform from spiking to fluorescence in GCaMP6s-expressing neurons in Camk2a-tTA;tetO-GCaMP6s mice. A) Top: Trial-averaged instantaneous spike rate of a neuron for trials that contain specific numbers of spikes (mean with s.e.m.). The neuron is recorded *in vivo* via two photon targeted loose patch during visual stimulation with oriented gratings. Bottom: Trial-averaged calcium responses of the same trials above (mean with s.e.m.). B) Average dF/F vs. spike rate per trial for the same example neuron. Gray lines indicate a linear regression with 95% C.I. C) As in B) but for a population of recorded cells (n=21). Box plots are shown for all trials that had a specific number of spikes. D) Histograms of the coefficient of determination (gray) and Pearson’s correlation (black) for all analyzed neurons.

**Fig. S8.**
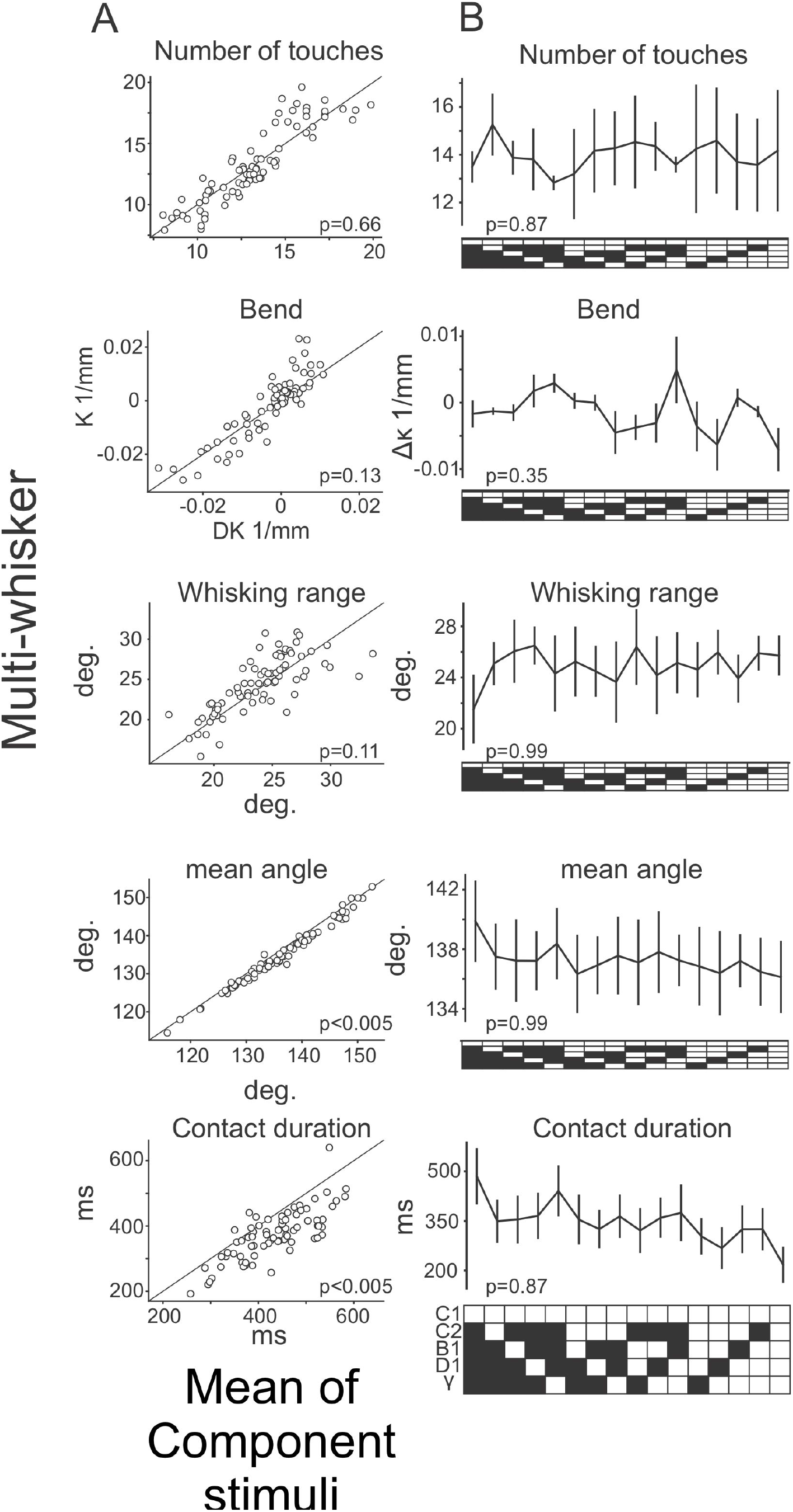
Whisker kinematics across the stimulus set. A) Scatter plots comparing the mean of each indicated kinematic variable during multi-whisker stimuli (y-axis) and the mean across the component single whisker stimuli for each corresponding multi-whisker stimulus (x-axis). B) The mean of each indicated kinematic variable for the C1 whisker compared across all stimuli that contain the C1 piston. N = 3 mice.

**Fig. S9.**
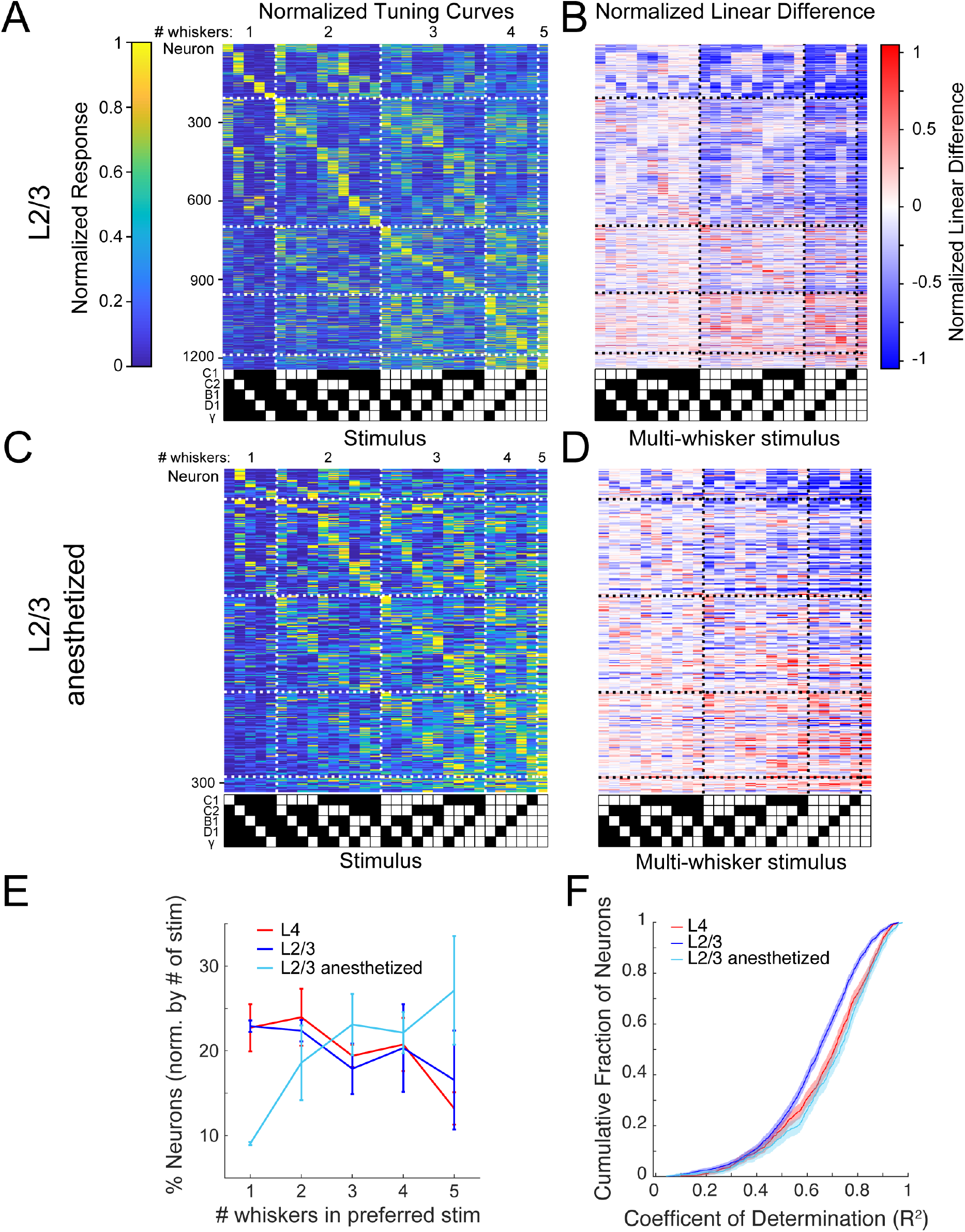
Altered population codes in anesthetized, passively stimulated mice. A) Normalized, cross-validated tuning curves for all sensory-driven L2/3 neurons from awake mice. The black diagonal line depicts the expected population diagonal if the neural population uniformly covers the stimulus space. B). Normalized linear difference for the neurons in A). C) Same as A) but for L2/3 neurons recorded in anesthetized mice whose whiskers were stimulated with piezoelectric benders. D). Normalized linear difference for the neurons in C). E) Percent of neurons whose preferred stimulus contains 1, 2, 3, 4, or 5 whiskers, normalized by the number of stimuli presented in each category (mean and 95% C.I.; L4 p = 0.10, n=3; L2/3 p = 0.72, n=3; L2/3 anesthetized p = 0.08, n=3; one-way ANOVA). F) Cumulative density functions of the coefficient of determination (R^2^) from a general linear model fit of tuning curves for L4, L2/3 awake, and L2/3 anesthetized neurons (L2/3 vs L2/3 anesthetized p < 10^−17^; L4 vs L2/3 p < 10^−9^; L4 vs L2/3 anesthetized p = 0.03, Wilcoxon ranksum, Bonferroni multiple comparisons correction; L4 n=847, L2/3 n=1415, L2/3 anesthetized n=635).

**Fig. S10.**
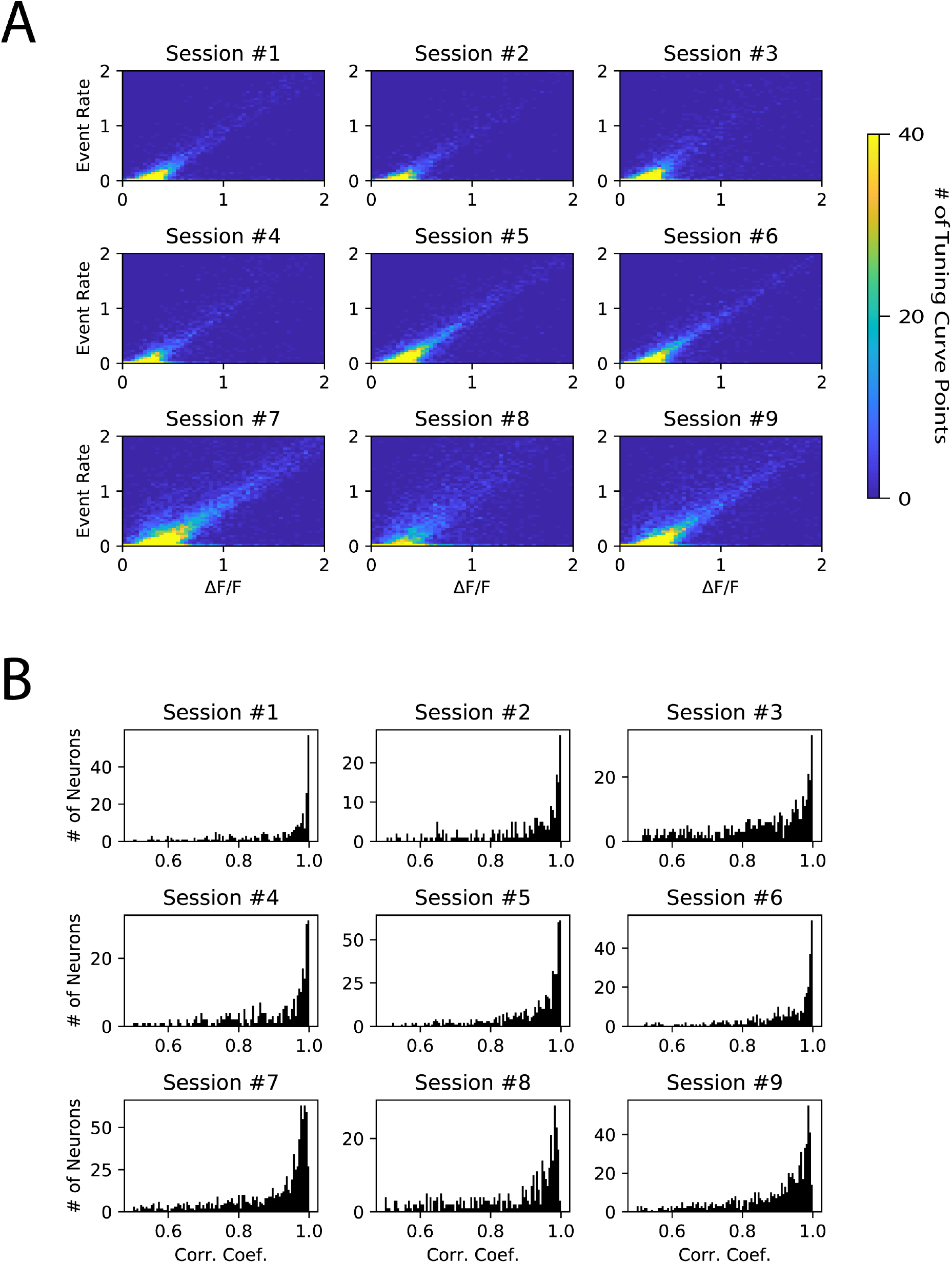
Effect of deconvolution on tuning curve calculation. A) 2-D histogram of tuning curve elements computed using ΔF/F as compared to deconvolved event rate. Each plot is from one imaging session (either S1 L2/3 or S1 L4). Only trials following catch trials were analyzed, to avoid contamination from decaying calcium transients from the previous trial. B) Histograms of correlation coefficients between tuning curve computed with deconvolved event rate, and tuning curve computed with ΔF/F. One plot per S1 imaging session.

**Fig. S11.**
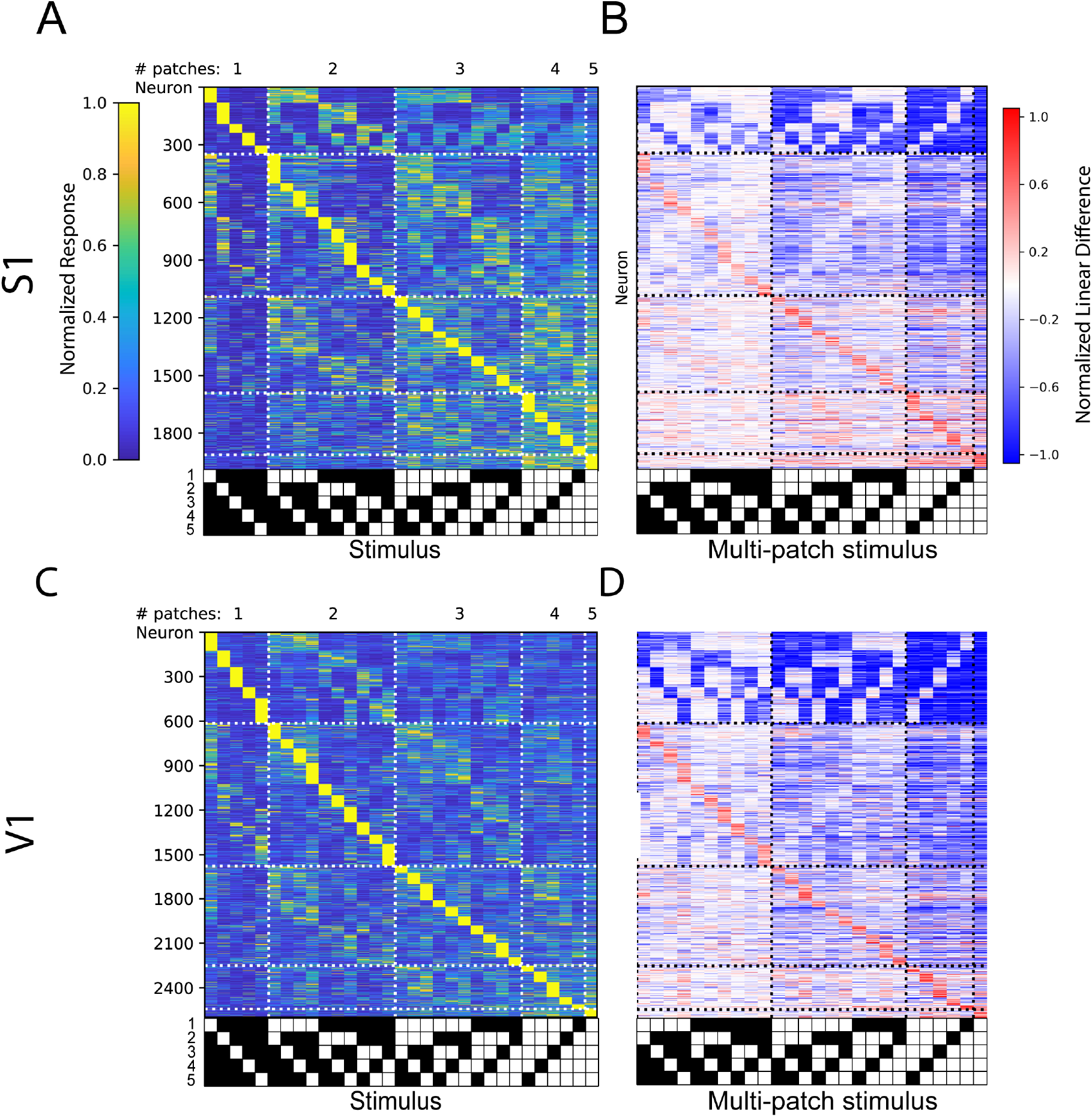
Non-cross validated tuning curves and linear difference plots of S1 and V1 neurons. A) Non-cross validated tuning curves and B) linear difference plots for S1 layer 4 and layer 2/3 experiments, sorted on and displaying average of all trials, rather than sorted based on held out trials. C,D) Same as A),B), for V1.

## Methods

### Experimental model details

All experiments were performed on mice between 1.5 to 9 months of age. CaMKII-tTA mice (RRID:IMSR_JAX:003010) crossed to tetO-GCaMP6s mice (RRID:IMSR_JAX:024742) were used when imaging L2/3. For S1 experiments, both lines had been outcrossed to the ICR line (Charles River) for several generations. Experiments imaging L4 in S1 used Scnn1a-Tg3-Cre mice (RRID:IMSR_JAX:009613) that had been outcrossed to the ICR line for several generations. For V1 L2/3 experiments, mice were on a mixed background between outcrossed tetO-GCaMP and camk2-tTA on the C57/B6 background. For V1 L4 experiments Scnn1a-Tg3-Cre mice were crossed to Ai162(TIT2L-GC6s-ICL-tTA2)-D mice (RRID:IMSR_JAX:031562). Both female and male animals were used and maintained on a 12:12 reversed light:dark cycle. All procedures were approved by the Animal Care and Use Committee of UC Berkeley.

### Preparation for *in vivo* two photon imaging

Headplate attachment, habituation to running on a circular treadmill, intrinsic imaging, and cranial window installation were performed as described previously (Pluta et al., 2017). Briefly, anesthesia was induced with 5% isoflurane and maintained at 1-3% during surgery. Respiratory rate and response to toe/tail pinching was monitored throughout surgery to ensure adequate anesthetic depth. 0.05 mg/kg of buprenorphine was administered subcutaneously for post-operative analgesia. The scalp was disinfected with 70% alcohol and 5% iodine. The skin and fascia above the sensory cortices were removed and Vetbond (3M) was applied to the skull surface and wound margins. A custom stainless steel headplate was fixed to the skull with dental cement (Metabond). For S1 mice, two days after surgery, mice were habituated to head-fixation on a free-spinning circular treadmill. This was repeated for 4-8 days until they freely ran at a fast and steady pace (>30 cm/s). Since mice often run in bouts, criterion was set such that mice had to maintain an average run speed above 10 cm/s over an hour to be included in the S1 component of the study.

Scnn1a-Tg3-Cre mice that reached running criterion were injected with AAV9-flexed-CAG-GCaMP6s (UPenn Vector Core) virus in left S1. Briefly, they were anesthetized and administered buprenorphine as described above. A dental drill (Foredom) was used to create a small bur hole 1.3 μm posterior and 3.5 μm lateral to bregma (marked previously with a sharpie during the headplate procedure). Then a WPI UltraMicroPump3 injector was used to inject 300 nL of the virus at a depth of 350 μm and a rate of 0.5 nL/s. Post-injection, the needle was left in the brain for 5 minutes to allow the viral solution to absorb into the tissue. Injected mice were provided 2-3 weeks with intermittent head-fixation over the circular treadmill to allow the infected neurons to ramp up expression of GCaMP6s.

A cranial window was installed to provide for optical access to the cortex. The mice were anesthetized and administered buprenorphine as described above, and administered 2 mg/kg of dexamethasone as an anti-inflammatory. Post-dexamethasone injection, isoflurane was maintained at 1-1.5% during surgery. A dental drill was used to drill a 3mm diameter craniotomy over the left primary somatosensory cortex. For V1 experiments, a biopsy punch was used to create a 3.5mm diameter craniotomy over the left primary visual cortex. A window plug consisting of two 3 mm diameter coverslips glued to the bottom of a single 5 mm diameter coverslip using Norland Optial Adhesive #71 was placed over the craniotomy and sealed permanently using Orthojet. Mice were provided at least two days to recover.

For S1 experiments, intrinsic optical imaging was performed through the cranial window to localize the C1 and C2 barrels. Prior to imaging, anesthesia was induced as described above and then the mice were administered 5 mg/kg Xylazine. Anesthesia was maintained with 1% isoflurane during imaging. Imaging and stimulation was conducted using custom software written in MATLAB. Mice were given 24 hours to recover.

### Tactile stimulus presentation and *in vivo* imaging of S1

Aluminum was custom machined to hold 5 pneumatic pistons on ball joints equally spaced along a circle. Pistons had a diameter of 1.6 mm, a length of 3 cm, and their tips were cut in such a way as to provide a flat surface for the whisker to palpate against. The apparatus was attached to a magnetic stand that could easily be slid along the optical table and locked in place.

Prior to the experiment the mouse’s whiskers were trimmed. Anesthesia was induced and maintained as described above. All whiskers were trimmed completely off, sparing the C2, C1, B1, D1, and Gamma whiskers. If one of these whiskers was missing, a nearby whisker was substituted (one mouse D2 for D1; one mouse beta for B1). The remaining 5 whiskers were trimmed in a staircase like fashion such that the tip of the anterior whiskers in the C row wouldn’t overlap with the tips of the posterior whiskers during whisking. Mice were head-fixed on a freely spinning running wheel under a Nixon 16x-magnification water immersion objective and imaged with a two-photon resonant scanning microscope (Neurolabware) within a light tight box. Each piston was extended one-by-one and adjusted such that its corresponding whisker contacted it close to the peak of the whisker’s protraction. A stereoscope and high-speed whisker tracking camera were used in positioning the piston and to verify contact was specific and repetitive. Calcium imaging occurred at 15.45 Hz with fields of view (FoVs) ranging from 800 μm by 1 mm to 1 mm by 1.3 mm. Wide-field reflectance imaging with a blue LED was used to illuminate the vasculature and center the FoV on the region the intrinsic signal identified as corresponding to the C1 barrel. For L2/3 imaging, imaging depth was 100 – 300 μm, and for L4 imaging, depth was 350 – 500 μm deep. In some experiments, 4 depths 33 μm apart were imaged sequentially using an electrotunable lens (Optotune), with each depth sampled at an effective frame rate of 3.86 Hz.

Piston combinations were presented sequentially to the mouse’s whiskers in a pseudo-random fashion where more weight was given to stimuli that had been presented the least. Stimulus presentation lasted one second followed by a two second inter-trial interval. Average running speed per trial was computed online, and trials where the mouse ran slower than 10 cm/s were repeated later in the experiment. The experiment finished once each of the 32 stimulus combinations (including a catch/no stimulus condition) was presented 20 times while the mouse was running.

### Visual stimulus presentation, *in vivo* imaging of V1, and pupil tracking

For visual stimulus presentation, the monitor was placed 14-15 cm from the eye. Animals were habituated to visual stimulation on the setup for at least two sessions prior to imaging. The visual stimulus subtended one of five contiguous square patches laid out in a cross pattern, to match the spatial layout of pistons in the tactile stimulation experiments. Because contiguous homogeneous textures have been widely reported to drive sublinear responses, we focused on heterogeneous textures, with surround patches shifted in direction by 90 degrees relative to the central patch. Trials with patches 10 visual degrees in diameter were interleaved with trials having patches 15 degrees in diameter, and in some experiments, trials with 5 degree patches. To match the spatial scale of single S1 barrels, only the analysis of 10 degree patch presentation is presented in the main figures. Within each patch, square wave drifting gratings of zero or ninety degree orientation were shown, with a spatial frequency of 0.08 cycles per degree, and a temporal frequency of 1 Hz. Stimulus presentation lasted one second followed by a one second inter-stimulus interval. Patch configurations, orientations, and sizes were pseudorandomly interleaved, and stimuli were generated and presented using the Psychophysics Toolbox (Brainard, 1997). Each distinct visual stimulus was displayed for ten repetitions.

For V1 imaging, the same Neurolabware setup was used as described above. The imaging FOV was 430 by 670 um, with four planes spaced 37.5 μm apart imaged sequentially, sampling each plane at an effective frame rate of 7.72 Hz. Electrical tape was applied between the objective and the mouse’s headplate to block monitor light from entering the microscope.

Running was monitored in the same way as described above, with trials where maximum absolute run velocity > 1 cm/sec classified as “moving” and < 1 cm/sec classified as “non-moving”. Analysis was restricted to “non-moving” trials.

Eye movements were imaged using a Basler Ace aCA1300-200um camera, with a hot mirror (Edmund Optics) placed between the eye and the monitor reflecting infrared light for eye imaging while transmitting visible light for visual stimulation. Infrared illumination was provided by the two photon imaging laser, transmitted through the pupils, as well as a panel of 850nm LEDs (CMVision). Pupil location and diameter were tracked using custom MATLAB code. When splitting the data into halves for cross-validated tuning curve and linear difference visualizations (see below), the halves were chosen to be balanced for pupil diameter and location.

### Anesthetized stimulus presentation

Preparation, trimming, and imaging conditions were consistent with above, however this cohort of mice were not habituated to run while head-fixed. As well, immediately after whisker trimming the anesthetized mice, they were injected intraperitoneally with 0.075mg chlorprothixene dissolved in saline. Ten minutes later the isoflurane was turned down from 2% to 0.5% and the mice were injected intraperitoneally with 40mg urethane dissolved in saline. Mice were then head-fixed under the resonant scanning two photon microscope, but this time over a feedback controlled heater (FHC). Up to 1% isoflurane was provided to maintain anesthetic depth, but was generally not necessary. A custom 3D printed whisker collar attached to a piezo plate bender (Noliac) was slid over each of the 5 whiskers. A cosine waveform designed to match the rate of angle change observed during natural touch was used to stimulate the whiskers (data not shown). The stimulus occurred at 16 Hz matching the average number of touches between the C1 whisker and its corresponding piston observed during one randomly selected experiment. Stimulus combinations were presented in a block randomized fashion, with each trial lasting 3 seconds, and the stimulus occurring for either 0.5 or 1 second. The experiment finished once each stimulus combination was presented 20 times.

### Calcium imaging analysis

For the S1 data, motion correction, ROI identification, and neuropil correction were calculated as described previously (Pluta et al., 2017). Briefly, two photon movies were corrected for brain motion using Scanbox’s sbxalign script (MATLAB, Mathworks) to correct for the rigid, 2D translation of individual frames. Regions of interest (ROIs) encompassing neurons were identified in a semi-automated manner using Scanbox’s sbxsegmentflood (MATLAB) which computes and thresholds the pixel-wise cross-correlation for all pixels within a 60 by 60 pixel window. The ROI’s signal (*R*_*i*_) was taken as the mean value across all pixels within and unique to that ROI. This signal is assumed to be a mixture of the cell’s actual fluorescence signal and a contaminating neuropil signal resulting from scattering producing off-target excitation, high illumination powers producing out of focus fluorescence, or unresolvable neurites passing through the microscope’s point spread function. The neuropil signal (*N*_*i*_) for each ROI was computed by averaging over an annulus of pixels surrounding the ROI but excluded pixels assigned to other ROIs as well as a smaller annulus of pixels that acted as a buffer in case any 2D motion artifact was not perfectly accounted for. This buffer annulus existed for all ROIs and was excluded from any neuropil calculation. As a result the max diameter of the neuropil annulus varied per ROI in order to ensure a similar number of usable pixels to average over. Each neuron’s neuropil-corrected fluorescence signal (*F*_*i*_) was computed per ROI by the following equation:

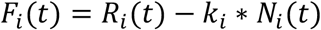

The amount of contamination (*k*_*i*_) was assumed to be constant per ROI, but vary between ROIs as a result of local differences in expression and scattering. Each *k*_*i*_ was defined by assuming that the neuron’s true fluorescence signal (*F*_*i*_) can never be negative (i.e. *k*_*i*_ * *N*_*i*_(*t*) ≤ *R*_*i*_(*t*)), and that the neuropil signal cannot contaminate more than its measured value. The contamination coefficient per neuron was defined as follows:

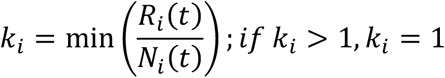

A baseline fluorescence 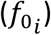 was calculated for each neuron by averaging its neuropil-corrected fluorescence over the last second of the inter-trial interval following a catch trial where no stimulus was presented, and then by averaging across the 20 catch trials presented. The change in fluorescence (ΔF/F) was calculated by subtracting off the baseline fluorescence from the neuropil-corrected fluorescence, and then dividing by the baseline fluorescence:

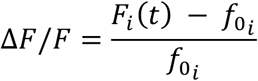

Each neuron’s response to a given trial was calculated as the mean ΔF/F over the one second stimulus period and subsequent two second inter-stimulus interval.

The C1 column was defined in a multistep process. First a pixel-wise average across all running trials for each stimulus was performed, followed by averaging across the one second stimulus period to define the mean response to each stimulus for the whole field of view. The mean response to the no stimulus (catch) condition was then subtracted off from all averages. Next these averages where multiplied by the stimulus matrix defining which whiskers were stimulated in each stimulus to generate a mean response profile for each whisker. Last each pixel was labeled by which whisker it responded the most strongly to creating a pixel-wise map of preferred whisker. A Gaussian smoothing operation was used as well as dilation and erosion steps to generate smooth blobs defining the location of each column. The non-C1 columns were ignored as the C1 column is the only one whose borders were well defined since it’s the only one whose surrounding whiskers were stimulated.

For the V1 data, motion correction and ROI segmentation was performed using Suite2p (Pachitariu et al., 2017). Neuropil subtraction was applied as described above. ΔF/F traces were calculated with baseline F_0_ computed over a sliding 20^th^ percentile filter of width 3000 frames. Because the inter-stimulus interval was reduced in V1 recordings to permit more stimuli to be displayed, calcium transients overlapped between successive trials. Therefore, we deconvolved calcium traces for this data using OASIS with L1 sparsity penalty (Friedrich et. al., 2017), using ΔF/F traces as input. We present a comparison of responses between deconvolved event rate and ΔF/F in Fig. S10, showing a tight correlation. Deconvolved event rates were normalized such that a train of events producing calcium transients with time averaged ΔF/F = 1, had a time averaged magnitude of 1.

In order to ensure that deconvolution of calcium responses in the V1 data did not introduce biases into the calculation of tuning curves, we compared tuning curves computed with ΔF/F with tuning curves computed using deconvolved event rates. For this comparison, we examined trials following non-stimulus trials in the S1 data, in which stimuli were sufficiently widely spaced in time that calcium transients did not overlap between trials. In this data, we found that tuning curve values computed using the two methods were highly correlated (Fig. S10a) with the vast majority of neurons showing Pearson correlation coefficients between deconvolved and ΔF/F tuning curves >0.9 in typical experiments (Fig. S10b).

### High-speed whisker tracking

In all trials the whiskers were imaged at high speed (300 frames per second) during the 1 second stimulus period using a camera (FLIR) with a telecentric lens (Edmund Optics) placed below the running wheel and reflected off a 45 degree mirror. The whiskers were illuminated in a trans-illumination fashion by a panel of 850nm LEDs (CMVision) covered with a piece of tracing paper acting as a diffuser. The illumination source was placed at a sufficient distance as to create a flat background of uniform intensity.

Whiskers were later identified and tracked across time in a subset of experiments using DeepLabCut(Mathis et al., 2018). The algorithm was trained on each mouse individually using ~160 frames spanning all piston conditions in the experiment. Frames in each condition were chosen automatically using k-means clustering (according to intensity values in pixel space; one frame was chosen from each cluster, k = 5). In each frame we labeled 30 points, 6 for each whisker (in equal distances), that were used as target values for the neural network. After evaluating whisker locations in all test videos, we fitted 2nd degree polynomial function for each whisker points using random sample consensus method (RANSAC; with 4 minimum samples for the fitting) deriving whisker trajectory in each frame. Whisker tracing outliers were discarded and replaced with the mean whisker trajectory from flanking frames. The estimated mechanical parameters: the whisker angle was estimated by fitting linear curve to the whisker origin and was defined relative to the frame vertical axis. Subsequently, 95% of the whisker angle histogram was defined as the whisking range in a trial. The whisker bend (*κ*) was calculated at the third lower part of the whisker as 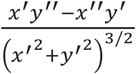 where x and y are the pixel coordinates of the whisker. To estimate the overall force applied to the whisker, we subtracted the free-whisking internal curvature from the curvature values in the trial. We defined touch as the first contact of the whisker with its matching piston in a series of successive contacts.

For each kinematic variable we compared the multi-whisker trials (e.g., trials in which C1, C2 and D1 are stimulated) and the single-whisker trials (C1, C2 and D1 individually). Each parameter of the stimulated whiskers was averaged across all trials with a specific piston combination and averaged also within a trial (for the multi-whisker trials) or between trials (for the single-whisker trials). We compared each kinematic variable using paired student’s t-test and corrected with the Benjamini & Hochberg procedure for multiple comparisons (*α* = 0.05). We further compared the kinematic variables for the C1 whisker in all conditions involving it using a 1-way ANOVA.

### Two photon targeted loose patch calibration of GCaMP6s signaling

Adult CamkII-tTa;tetO-GCaMP6s mice were anesthetized with 2% isoflurane and kept warm with a feedback controlled heater (FHC). After scalp and periosteum removal a drop of Vetbond was applied to the skull, and a small custom headplate was affixed to the skull with Metabond dental cement. A 3 mm circular craniotomy over left primary visual cortex was performed with a biopsy punch, and the brain was then covered with warm 1.2% agarose in PBS. The dura was typically left intact. The mouse was injected with chlorprothixene (5 mg/kg) and urethane (1 g/kg). The mouse was mounted under a Sutter MOM two photon microscope equipped with a 20x 1.0 NA objective (Olympus) operated with ScanImage software (Vidrio). Anesthesia was maintained in a stable state with the addition of 0-1% isoflurane. Patch pipettes were filled with standard ACSF contained 50μM alexafluor 594 K+ salt (Thermofisher). The pipette was advanced under positive pressure and two photon guidance through the dura/pia and towards GCaMP6s-expressing neurons in L2/3 at a 27 degree angle, carefully avoiding any blood vessels. During pipette advancement full field high contrast drifting gratings were presented to generate neuronal responses in order to identify responding neurons by visual inspection. Cells with strong responses were targeted with the electrode in loose patch configuration using a Multiclamp 700B amplifier (Molecular Devices). Following acquisition of a low resistance seal, negative pressure was applied to increase the recorded amplitude of action potentials so that they were substantially larger than the background electrical noise, facilitating spike detection. The experiments then commenced. Trials were 3 seconds in length and consisted of a 500 ms baseline (grey screen) after which a high contrast grating drifted at 4-8 directions for 2.5 seconds, with 2 second intertrial intervals (grey screen). Visual stimuli were generated with Psychophysics toolbox(Brainard, 1997) and were displayed on 7” LCD monitor (60 Hz refresh rate), positioned ~5-7 cm from the contralateral eye. All visual stimuli were full screen (~40 degrees of visual angle) square wave drifting gratings with a temporal frequency of 2 Hz and a spatial frequency of 0.04 cycles per degree at 100% contrast. The LED backlight of the monitor was controlled by a high speed LED driver (Mightex) and triggered by the turn around signal from resonant scan mirror via a custom Arduino-based circuit so that the monitor was only illuminated during mirror flyback and not when the PMTs were being sampled to avoid any optical artifacts from the monitor. A trigger from the visual stimulus computer synchronized the electrophysiology and two photon imaging computers. Imaging power was 50-75 mW at the exit of the microscope objective. Electrophysiology data was acquired at 20 kHz and sampled via a National Instruments card (PCIe-6323) via custom MATLAB scripts. Spikes were extracted after digital high pass filtering (>1 kHz) and detected by a threshold set as samples greater than 6 times the standard deviation of the noise. Imaging data was analyzed as above following motion correction (which was minimal, but corrected nonetheless).

### Quantification and statistical analysis

Statistically significant differences between conditions were determined using standard parametric and nonparametric tests in MATLAB, including a 1-way ANOVA, 2-way ANOVA, two-sample t-test, Wilcoxon rank sum test, and a Kolmogrov-Smirnov test. Analyses were performed on each ROI’s measured ΔF/F for each trial. A single trial’s response was calculated as the average ΔF/F during the entire 1 second of stimulation. Analysis was limited to ROIs that were significantly driven by at least one stimulus. A significant response for a stimulus had to meet two criteria: have a mean ΔF/F greater than 0, and pass a two-sample t-test between the evoked responses for a given stimulus and the measured ΔF/F values during catch trials. The Benjamini & Hochberg false discovery rate correction was used to correct for the multiple comparisons taken across the multiple stimuli. Outlier responses per stimulus condition were identified by the median rule, where values further than 2.3 times the inter-quartile range from the median are determined to be outliers, and were removed prior to any analysis. Tuning curves were generated by averaging across all inlier trials, and 95% confidence were generated via bootstrap. In the awake mouse, these trials were further limited to running trials defined as trials where the mouse’s run speed was above 10 cm/s and were not outliers as determined by again applying the median rule. Tuning curves subtracted off the mean response to catch trials to correct for any tuning offset.

The principal whisker was determined by fitting a general linear model to the individual trial responses as a function of what whiskers were stimulated. The model had one term for each whisker as well as an offset term.

A neuron was labeled as a single whisker neuron if it only exhibited a significant response to one of the five single whisker stimuli. It was labeled as a multi-whisker neuron if it responded significantly to two or more of the five single whisker stimuli. Or it was labeled as a combination neuron if it wasn’t significantly driven by any single whisker stimuli, but was significantly driven by a multi-whisker stimulus.

For analyses comparing neurons’ functional responses to their barrel column, barrel columns were first defined via a custom segmentation algorithm applied to a pixel-wise piston preference map. First, a pixel-wise average response was calculated for each piston by averaging over all trials in which that piston was presented. Then a 2D Gaussian smoothing filter was applied to each average image to reduce noise. Next a pixel-wise preference map was created via labeling each pixel by the piston that produced the largest average response. Some pixels lie within resolvable neurons, however most pixels lie between neurons and reflect the neuropil signal, an average of the surrounding population of neurons and fibers of passage. This preference map was then segmented into five discrete, non-overlapping blobs following an erosion, dilation, erosion, and Savitzky-Golay filter steps to ensure smooth, column-like edges (MATLAB). The segmentation process was iterated over to ensure the identified barrel columns were consistent with the known somatotopic map and architecture. Neurons were assigned to a barrel column if their centroid lied within an identified column.

In order to ensure robustness of measured tuning properties, data for population-wide tuning curve and linear difference visualization was split into two halves, with trials balanced for stimulus conditions as well as pupil diameter and location in the case of the V1 data. Analysis was restricted to neurons showing a Pearson correlation coefficient >0.5 between tuning curves computed on the two halves of the data (i.e., neurons whose tuning curves could be reliably estimated based on half of the data). One half of the data, the “training set,” was used to estimate stimulus preference, used for sorting the neurons. The other half of the data, the “test set,” was displayed. In this way, the “preferred stimulus” was not mathematically constrained to evoke the largest response in the test set, and could be taken as a robust feature of the data where this was the case. A similar procedure was used to produce the population-averaged sorted linear difference plots. Tuning plots for the full data set are presented in Fig. S11.

## Data Availability

The data that support the findings of this study are available from the corresponding author upon reasonable request.

## Acknowledgements

The authors acknowledge the GENIE Project, Janelia Farm Research Campus, and the Howard Hughes Medical Institute for the GCaMP6 viruses, as well as Dan Feldman and members of the Adesnik and Feldman labs for comments on the manuscript, and technical support from Kiarash Shamardani. H.A. is a New York Stem Cell Foundation-Robertson Investigator. This work was supported by The New York Stem Cell Foundation. This work was supported by NINDS grant DP2NS087725-01, NEI grant R01EY023756-01.

